# Predicting multimodal MRI outcomes in children with neurodevelopmental conditions following MRI simulator training

**DOI:** 10.1101/2021.01.28.428697

**Authors:** Anish K. Simhal, José O. A. Filho, Patricia Segura, Jessica Cloud, Eva Petkova, Richard Gallagher, F. Xavier Castellanos, Stan Colcombe, Michael P. Milham, Adriana Di Martino

## Abstract

Pediatric brain imaging holds significant promise for understanding neurodevelopment. However, the requirement to remain still inside a noisy, enclosed scanner remains a challenge. Verbal or visual descriptions of the process, and/or practice in MRI simulators are the norm in preparing children. Yet, the factors predictive of successfully obtaining neuroimaging data remain unclear. We examined data from 250 children (6-12 years, 197 males) with autism and/or attention-deficit/hyperactivity disorder. Children completed systematic MRI simulator training aimed to habituate to the scanner environment and minimize head motion. An MRI session comprised multiple structural, resting-state, task and diffusion scans. Of the 201 children passing simulator training and attempting scanning, nearly all (94%) successfully completed the first structural scan in the sequence, and 88% also completed the following resting state fMRI scan. The number of successful scans decreased as the sequence progressed. Multivariate analyses revealed that age was the strongest predictor of successful scans in the session, with younger children having lower success rates. After age, sensorimotor atypicalities contributed most to prediction. Results provide insights on factors to consider in designing pediatric brain imaging protocols.

## 1. Introduction

Pediatric brain imaging has made significant advances in non-invasively capturing *in vivo* the brain organization in typical and atypical youth using MRI (Oldehinkel et al. 2013; Craddock et al. 2013; Di Martino et al. 2014). Although promising, progress remains challenged by artifacts, most notably, head motion. Indeed even submillimeter head motion has been shown to introduce false findings that can affect between-group analyses and replicability (Yuan et al. 2009; Power et al. 2012; Satterthwaite et al. 2013; Yan et al. 2013; Yendiki et al. 2014; Zuo et al. 2014; Alexander et al. 2017; Oldham et al. 2020). Such artifacts are particularly notable for children with neurodevelopmental conditions, such as autism spectrum disorder (ASD) or attention-deficit/hyperactivity disorder (ADHD) (Yerys et al. 2009). As a result, continued progress depends on the need to collect high quality imaging data, which can only be obtained when children keep their heads virtually motionless when being scanned.

Multiple methods for addressing motion artifacts post-scan exist, but they inevitably limit both data and sample size, and thus degrees of freedom (Yan et al. 2013; Bright, Tench, and Murphy 2017; Ciric et al. 2017; Satterthwaite et al. 2019; Eklund et al. 2020). As a result, the prevailing wisdom remains - the best way to handle motion is to prevent it (Ai et al. 2020). In this regard, efforts to minimize motion during MRI scanning such as passive movie viewing (Vanderwal et al. 2015), real-time motion monitoring and/or feedback (Dosenbach et al. 2017; Greene et al. 2018; Krause et al. 2019), prospective motion correction (Ai et al. 2020), and head stabilizers (Power et al. 2019) have been reported to be effective. However, they may not all apply across the broad range of MRI modalities and specialized sequences, the list of which continues to emerge (e.g., fMRI, diffusion MRI, MR spectroscopy, quantitative T1-weighted/T2 mapping, arterial spin labeling). Solutions that can impact the broad range of brain imaging modalities are needed, as studies increasingly seek to obtain multiple structural and functional metrics to support biomarker discovery and/or delineate pathophysiological mechanisms. Similarly, in clinical settings, multimodal imaging is commonly used to increase diagnostic precision and guide treatment such as pre-surgical MRI for epilepsy or tumor removals (Jung and Lee 2010). Additionally, the tolerability and utility of emerging motion prevention approaches during scan sessions for more challenging populations, such as those with neurodevelopmental conditions, have yet to be comprehensively established.

To bypass these challenges, preparing children before scanning remains a critical requisite for pediatric brain imaging. Numerous studies have shown that preparation improves compliance and reduces anxiety related to the unfamiliar MRI scan environment (Gabrielsen et al. 2018; Ashmore et al. 2019). Preparation protocols have included showing child friendly books or videos (Barnea-Goraly et al. 2014), playing with MRI toys (Cavarocchi et al. 2019), immersion in virtual reality (Ashmore et al. 2019; Garcia-Palacios et al. 2007), or practicing in an MRI simulator ort “mock scanner”. MRI simulators are widely used and, unlike most other preparation methods, also allow direct training for motion control. As summarized in Table 1, to date, 12 studies using MRI scan simulator training have reported their utility in obtaining good quality data in one or two MRI brain scans collecting different MRI modalities in one session (Rosenberg et al. 1997; Epstein et al. 2007; De Bie et al. 2010; Barnea-Goraly et al. 2014; Theys, Wouters, and Ghesquière 2014; Nordahl et al. 2016; Gabrielsen et al. 2018; Thieba et al. 2018; Sandbank and Cascio 2019; Horien et al. 2020; Pua et al. 2020; Yamada et al. 2020). However, the factors predicting successful completion of multiple scans in a MRI session remain largely unknown. Understanding the specific role of predictive features can guide the development of child-specific protocols for MRI data collection as well as MRI simulation training. This will be particularly relevant for children with neurodevelopmental conditions.

**Table 1.**
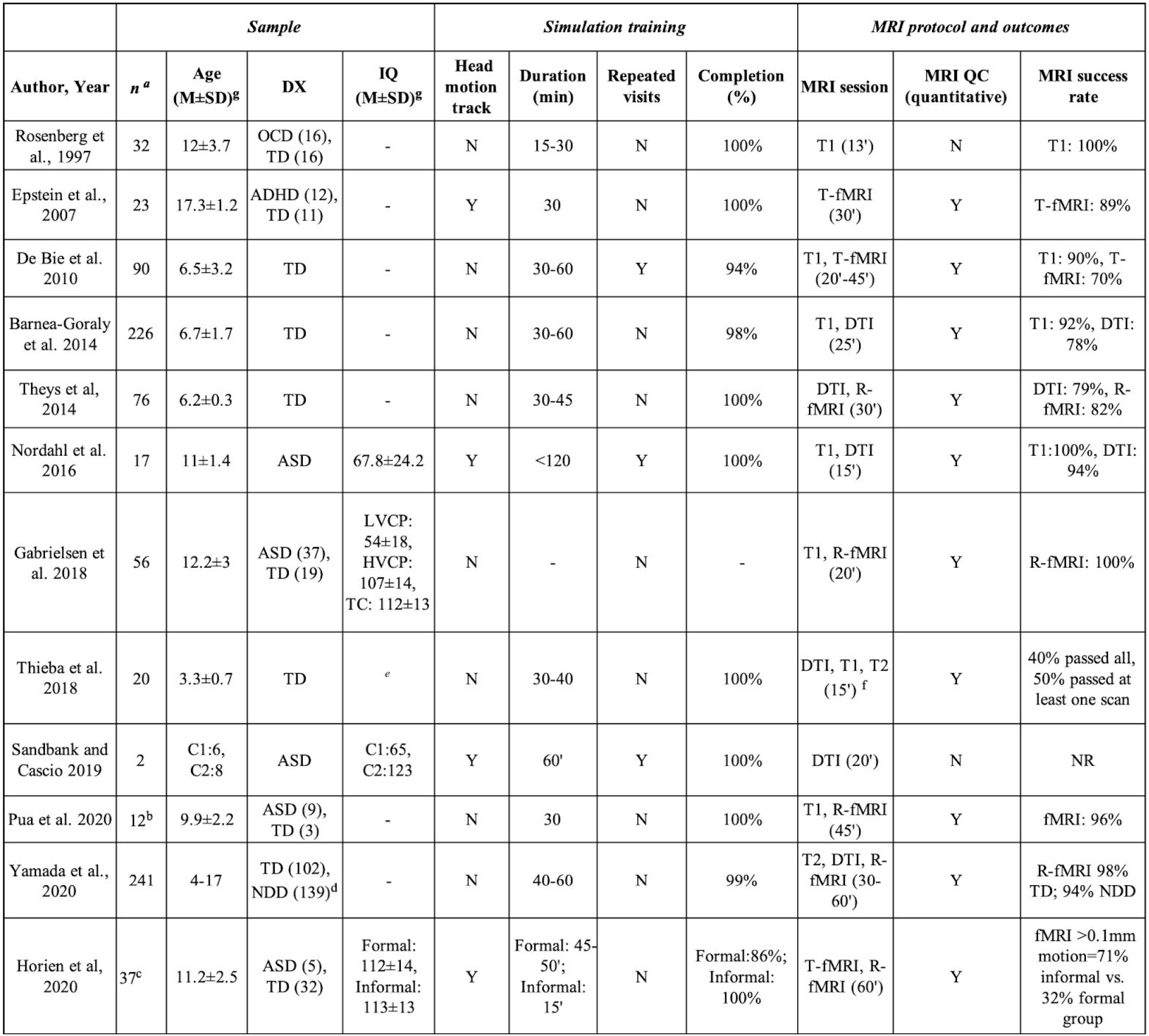
Overview of MRI simulator training studies in children. ^a^ Number of children attempting MRI simulator training protocol. ^b^ n=6 pairs of monozygotic twins concordant (3 pairs) or discordant (3 pairs) for ASD. ◻ Included a sample of 21 children (n=14 with a formal MRI simulator training, 7 without) and another sample of 16 children (all undergoing a formal training and used as a replication sample). ^d^ Included congenital genetic syndromes, ADHD, ASD, mild intellectual disability with unknown etiology, other behavioral and developmental disorders. ^e^ Used the Developmental NEuroPSYchological Assessment (NEPSY) (Korkman et al. 2007) and the Bayley Scales of Infant Development instead of standard IQ scores (Lennon et al. 2008). ◻ Resting-state fMRI (R-fMRI) was also acquired as time permitted but not analyzed. ^g^Range is provided for studies not reporting mean (M) and standard deviation (SD) and M and SD cannot be derived. Abbreviations: ADHD, attention deficit hyperactivity disorder; ASD, autism spectrum disorder; DD, developmental disability, DX, diagnosis; IQ, intelligence quotient, *n*, number of subjects; N, no; TD, typically developing; Y, yes.

Most studies using MRI simulator training have not characterized their samples in regard to behavioral, clinical or cognitive features. Of three notable exceptions (Nordahl et al. 2016; Thieba et al. 2018; Sandbank and Cascio 2019), only one examined the relation between children’s characteristics and MRI success (Thieba et al. 2018). Specifically, in 20 typically developing preschoolers children completing three structural MRI scans Thieba and colleagues found that children with higher language and cognitive skills were more likely to have successful scans following training (Thieba et al. 2018). Whether these findings extend to a wider age range and to children with neurodevelopmental conditions remains unexamined. Further, while 62% of the neuroimaging studies reviewed in Table 1 focused on either ASD or ADHD and/or other neurodevelopmental conditions, none has examined children with ASD and with ADHD, despite accumulating evidence of their frequent co-occurrence (Reiersen and Todd 2008; Simonoff et al. 2008; Rommelse et al. 2010; Grzadzinski et al. 2011; Leitner 2014; Kern et al. 2015; Joshi et al. 2017).

With these considerations in mind, here we report our effort to assess the role of a range of symptom domains in predicting completion of multimodal imaging data in N=250 verbally fluent school-age children with ASD and/or ADHD. Besides child characteristics, we also examined the contribution of MRI simulator training performance. MRI simulator training protocols have varied in terms of equipment, duration, frequency of the training sessions, as well as specific objectives. Specifically, while all studies aimed to acclimate children to the MRI environment by using in-house or commercial MRI simulators, some explicitly included training to decrease in-scanner motion (Nordahl et al. 2016; Sandbank and Cascio 2019; Horien et al. 2020). Such motion training has been accomplished using verbal or visual feedback following qualitative direct observation or based on quantitative data from motion sensors. To date, only three studies used motion sensors to train either typically developing or children with ASD, albeit in small samples (n=2-19) (Nordahl et al. 2016; Sandbank and Cascio 2019; Horien et al. 2020). Thus, along with the child’s clinical characteristics, the present study assessed to what degree motion during the simulator training would contribute in predicting successful MRI data collection.

## 2. Methods

### 2.1 Participants

We examined data from 250 children aged 5.5 to 11.9 years participating in an ongoing study (NIH R01MH105506) of the neurobiological underpinnings of autistic traits in ADHD and/or ASD. Sample recruitment and characterization are detailed in the Supplementary Methods and in (Guttentag et al. 2021). During the course of the study, the enrollment and behavioral assessment site was transferred from the NYU Child Study Center, NYU Grossman School of Medicine, to the Child Mind Institute (CMI) when the principal investigator (ADM) moved to a new position. As detailed below, no demographic, nor clinical differences were noted across sites (Supplementary Table 1); nevertheless potential batch effects were addressed using the Bayesian method combating batch effects, ComBat (Johnson, Li, and Rabinovic 2006; Fortin et al. 2017). The study protocol was approved by the institutional review boards of NYU Grossman School of Medicine, and Advarra, Inc at CMI. Written parent informed consent and verbal assent were obtained for all participants and written assent was also collected for children older than seven years. All data were collected prior to the COVID-19 pandemic.

### 2.2 MRI simulator training protocol

The MRI simulator training session aimed to familiarize participants with the MRI scanning environment and protocol while training them to minimize head motion in the MRI simulator environment. To this end, we used an MRI simulator, a Siemens-32-channel mock head coil with mirror to see a screen on the back of the bore used to project the visual stimuli, a head motion tracking system, and the corresponding software package, all acquired from Psychology Software Tools Inc. (Sharpsburg, PA). The head motion tracking system relied on sensor hardware based on the Ascension Technology Corporation (now part of Northern Digital Inc., Waterloo, ON) Flock of Birds real time motion tracking system. Its accuracy is specified as 1.8mm root-mean-square (RMS) and 0.5° RMS; we confirmed spatial resolution was approximately 0.77 mm and 0.2° at a sampling rate of approximately 9 Hz. Similar MRI simulator equipment from the same vendor was used at both enrolling sites.

For all participants, a MRI simulator training session occurred at the first in-person diagnostic visit. The training consisted of five increasingly demanding steps during which children were asked to keep their head still. As illustrated in Figure 1, the training protocol began with a review of two social stories with text and pictures (Gray 2000). The first one described the MRI scan environment and requirements; the second one described the MRI simulator training. This first step was followed by the child laying on the bed of the MRI simulator while listening to 30 seconds of scanner gradient noises corresponding to the multimodal MRI sequences used in the real MRI session. This allowed the child to begin acclimation to the mock scan environment and scanner noise. Then, participants wore the motion sensor with a band on their forehead, while the mock Siemens 32 channel head coil and mirror were positioned. Afterwards, when the child agreed to do so, the table was slowly moved inside the simulator bore. Once inside the MRI simulator, children practiced controlling head motion for increasing durations of time using different audio visual stimuli and feedback. Each step was increasingly similar to a real MRI session by either increasing the time children were asked to stay still (from 2 to 6 minutes), and/or by removing the target visual feedback used in response to forehead movements greater than 1.5mm, the minimum motion detectable by the sensor. During the first and least demanding step, children completed a target game for two minutes. During that game they were asked to keep a white dot, representing the position of their forehead, in the center of a target projected on the screen. During the last two and most realistic steps, children watched a black screen with a centered white cross, reproducing the stimulus used for the R-fMRI scan in the real MRI session. The child moved from one step to the next when limited or no motion events > 1.5 mm were detected in a given task. Otherwise, each step was repeated until motion events > 1.5 mm ceased or became minimal. Once each of the five steps was completed successfully, a piece of a virtual puzzle projected in the screen was awarded. Upon completion of the whole virtual puzzle, children chose a toy from a box of rewards. A complete training session lasted approximately 30-60 minutes, including breaks as needed. If the child was unable to complete the training protocol in the first session, they were invited back until they successfully completed it. Only children who successfully completed a full MRI simulator training protocol were invited to a real MRI session.

**Figure 1.**
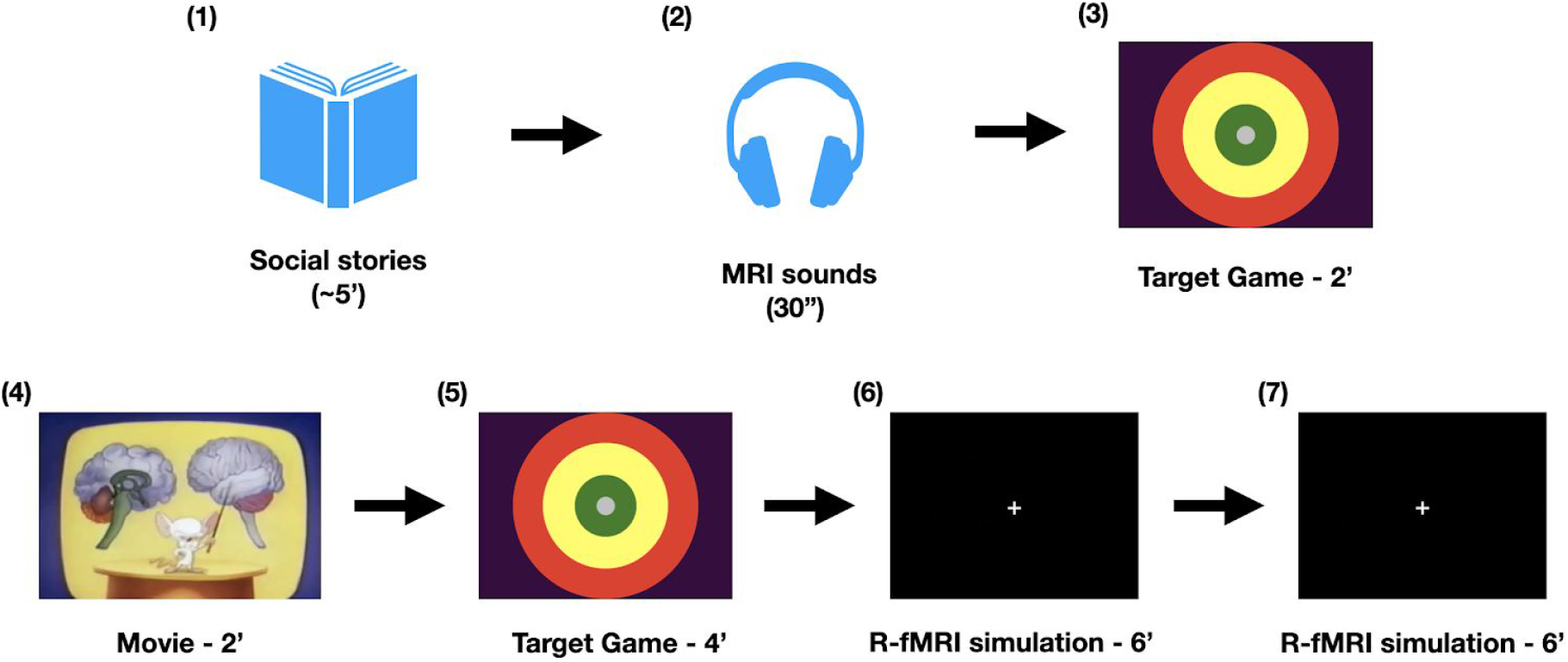
Overview of MRI simulation training protocol. First, the examiner and participant reviewed two “social stories,” one about the upcoming MRI session, the other about the “mock scan” training. Second, the child listened to 30-second-long multimodal MRI sounds outside the MRI simulator. For the 2 and 4 minute steps, the child played the target game inside the simulator during which children were instructed to keep a white dot (representing the position of their forehead indexed by the motion tracker) in the green center of the target. Between the two target game steps, a 2-min musical movie was played, stopping when head motion exceeded 1.5mm. Two 6-minutes blocks of R-fMRI simulations followed.

### 2.3 MRI protocol

All MRI images were collected at the NYU Center for Brain Imaging on a 3T Siemens Prisma scanner with the Siemens 32-channel head coil (Siemens, Erlangen, Germany). The study utilized a multimodal imaging protocol consisting of structural T1-weighted (T1w), T2 weighted (T2w), functional (rest and task), and diffusion MRI scans (see Box 1 for definition of most used MRI scanning terms in the manuscript). MRI scan parameters are detailed in Supplementary Table 2. The MRI session followed the same order of scan administration always starting with a T1-weighted scan, followed by a set of functional scans alternating rest and task scans, a T2w and a DTI scan completed the session (Figure 2). Head motion during the structural and diffusion scans was visually monitored through the operator window and via eye tracker camera positioned inside the MRI bore. During the functional scans, real time motion monitoring was also used (see Supplementary Methods). Administering the next scans in the sequence depended on completing the prior scan without notable motion; as such scans were repeated as needed. Children unable to complete all or some of the MRI scans in the first session were invited back for ‘make-up’ sessions, when possible. For consistency, the data analyzed here are based on the first MRI session.

**Table 2.**
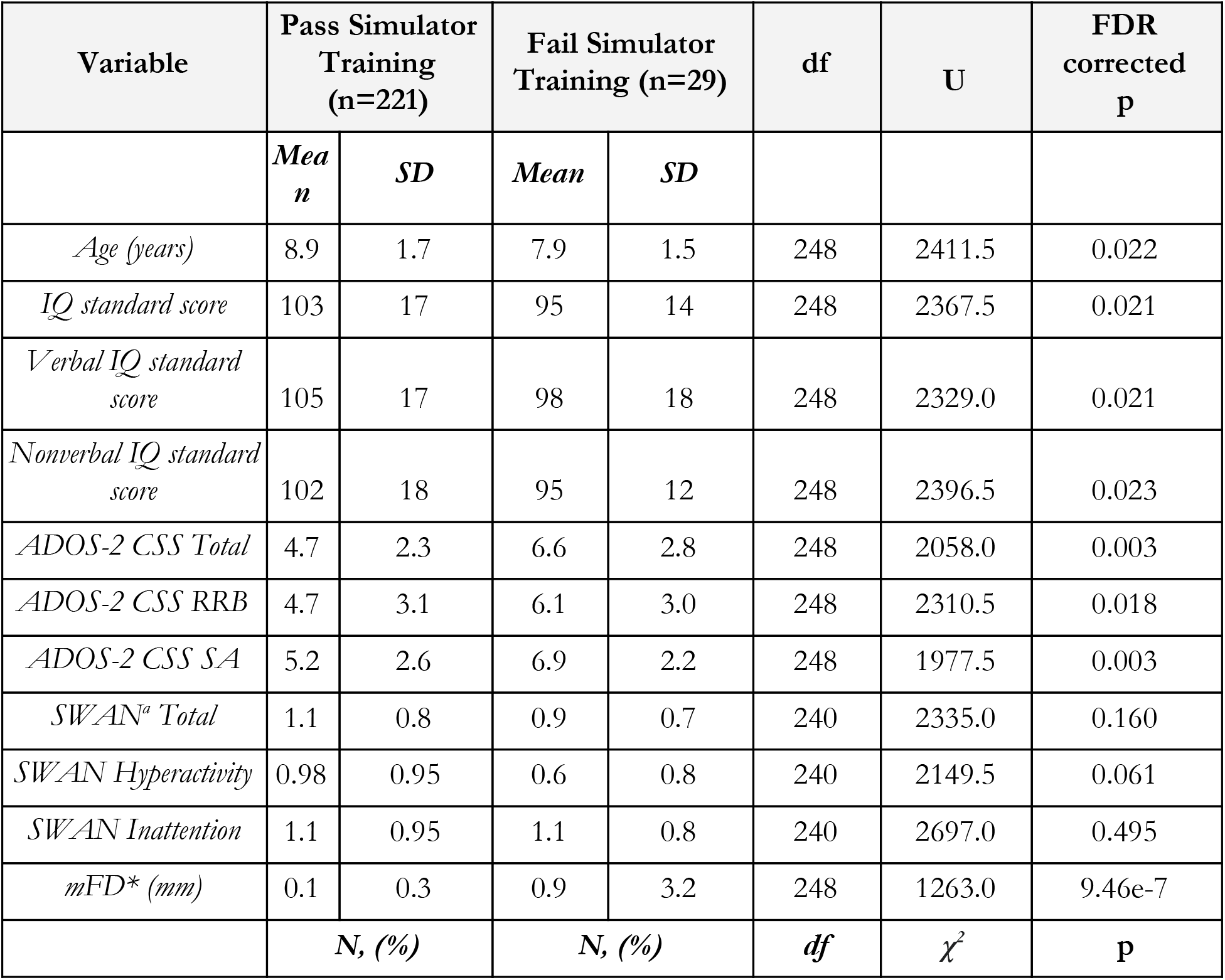
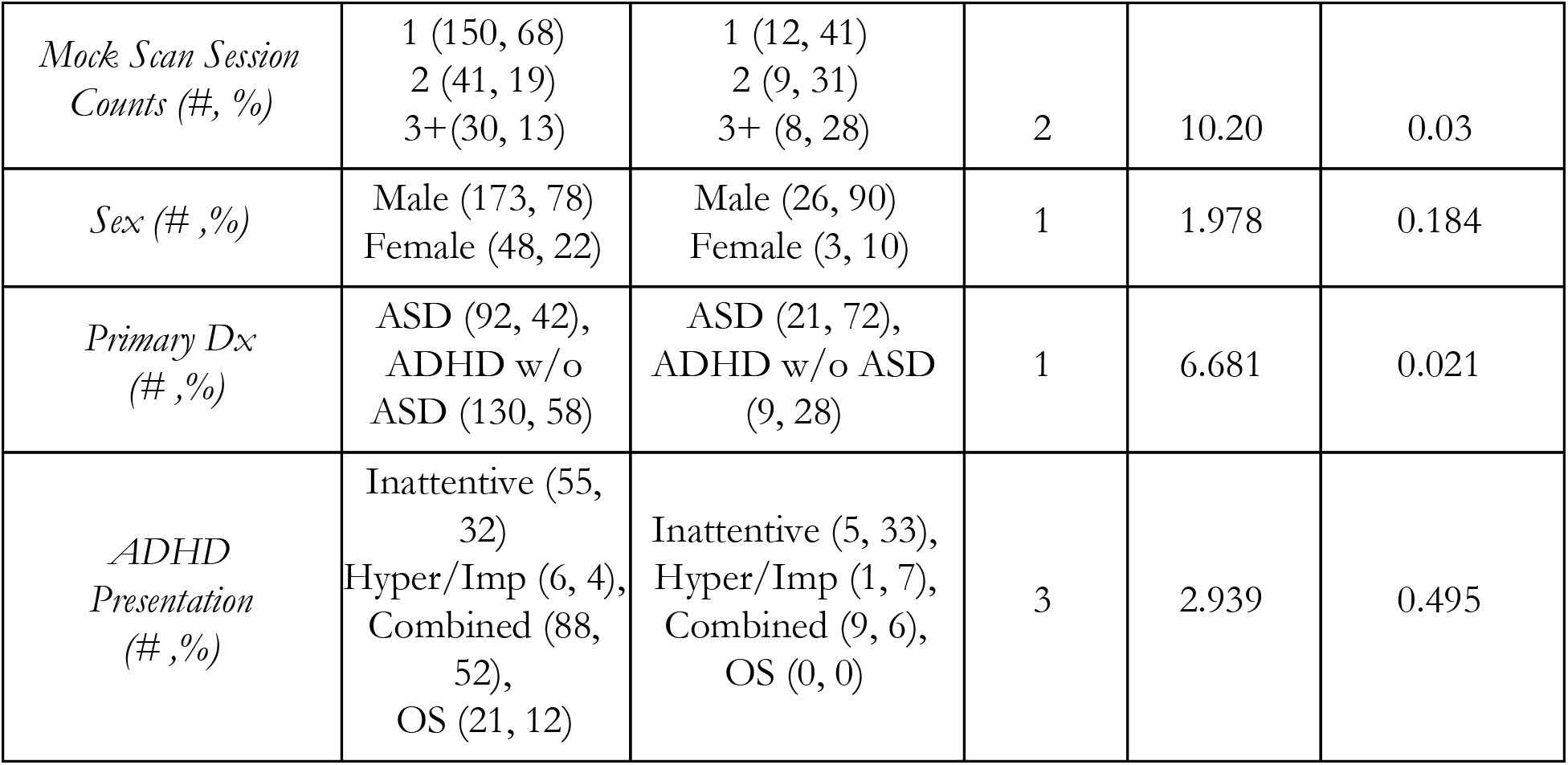
MRI simulator training outcomes. Group comparisons via Mann-Whitney U and Chi-square tests for continuous and categorical variables, respectively. All comparisons were corrected for multiple comparisons via false discovery rate - Benjamini-Hochberg (FDR-BH). ^a^ 12 children (8 passing and 4 failing the MRI simulator training) had missing SWAN parent scores. Abbreviations: ADHD, attention-deficit/hyperactivity disorder; ADOS-2, Autism Diagnostic Observation Schedule, second edition; ASD, autism spectrum disorder; CSS, calibrated severity scores; df, degree of freedom; Dx, diagnosis; mFD, mean framewise displacement (Jenkinson et al. 2002) data from the MRI simulator session; OS, otherwise specified; RRB, restricted and repetitive behaviors; SA, social affect; SD, standard deviation;, SWAN, Strengths and weaknesses of attention-deficit/hyperactivity symptoms and normal behaviors (average scores).

| Variable | Pass Simulator Training (n=221) |  | Fail Simulator Training (n=29) |  | df | U | FDR corrected p |
| --- | --- | --- | --- | --- | --- | --- | --- |
|  | *Mean* | *SD* | *Mean* | *SD* |  |  |  |
| *Age (years)* | 8.9 | 1.7 | 7.9 | 1.5 | 248 | 2411.5 | 0.022 |
| *IQ standard score* | 103 | 17 | 95 | 14 | 248 | 2367.5 | 0.021 |
| *Verbal IQ standard score* | 105 | 17 | 98 | 18 | 248 | 2329.0 | 0.021 |
| *Nonverbal IQ standard score* | 102 | 18 | 95 | 12 | 248 | 2396.5 | 0.023 |
| *ADOS-2 CSS Total* | 4.7 | 2.3 | 6.6 | 2.8 | 248 | 2058.0 | 0.003 |
| *ADOS-2 CSS RRB* | 4.7 | 3.1 | 6.1 | 3.0 | 248 | 2310.5 | 0.018 |
| *ADOS-2 CSS SA* | 5.2 | 2.6 | 6.9 | 2.2 | 248 | 1977.5 | 0.003 |
| *SWAN<sup>a</sup> Total* | 1.1 | 0.8 | 0.9 | 0.7 | 240 | 2335.0 | 0.160 |
| *SWAN Hyperactivity* | 0.98 | 0.95 | 0.6 | 0.8 | 240 | 2149.5 | 0.061 |
| *SWAN Inattention* | 1.1 | 0.95 | 1.1 | 0.8 | 240 | 2697.0 | 0.495 |
| *mFD\* (mm)* | 0.1 | 0.3 | 0.9 | 3.2 | 248 | 1263.0 | 9.46e-7 |
|  | **N, (%)** |  | **N, (%)** |  | **df** | **$\chi^2$** | **p** |

**Table 2. MRI simulator training outcomes.**
|  |  |  |  |  |  |
| --- | --- | --- | --- | --- | --- |
| *Mock Scan Session*<br>*Counts (#, %)* | 1 (150, 68)<br>2 (41, 19)<br>3+(30, 13) | 1 (12, 41)<br>2 (9, 31)<br>3+ (8, 28) | 2 | 10.20 | 0.03 |
| *Sex (#, %)* | Male (173, 78)<br>Female (48, 22) | Male (26, 90)<br>Female (3, 10) | 1 | 1.978 | 0.184 |
| *Primary Dx*<br>*(#, %)* | ASD (92, 42),<br>ADHD w/o<br>ASD (130, 58) | ASD (21, 72),<br>ADHD w/o ASD<br>(9, 28) | 1 | 6.681 | 0.021 |
| *ADHD*<br>*Presentation*<br>*(#, %)* | Inattentive (55,<br>32)<br>Hyper/Imp (6, 4),<br>Combined (88,<br>52),<br>OS (21, 12) | Inattentive (5, 33),<br>Hyper/Imp (1, 7),<br>Combined (9, 6),<br>OS (0, 0) | 3 | 2.939 | 0.495 |

**Figure 2.**
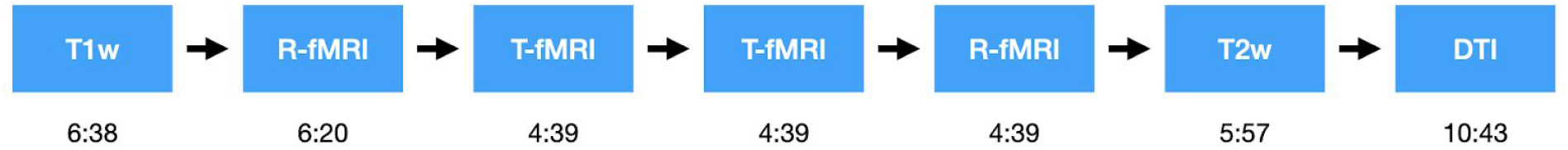
Scan sequence. Time is described in minutes:seconds. Moving to the next scan along the fixed order was dependent on the completion of the prior, scans were repeated as needed. See Supplementary Methods.

**Box 1.**
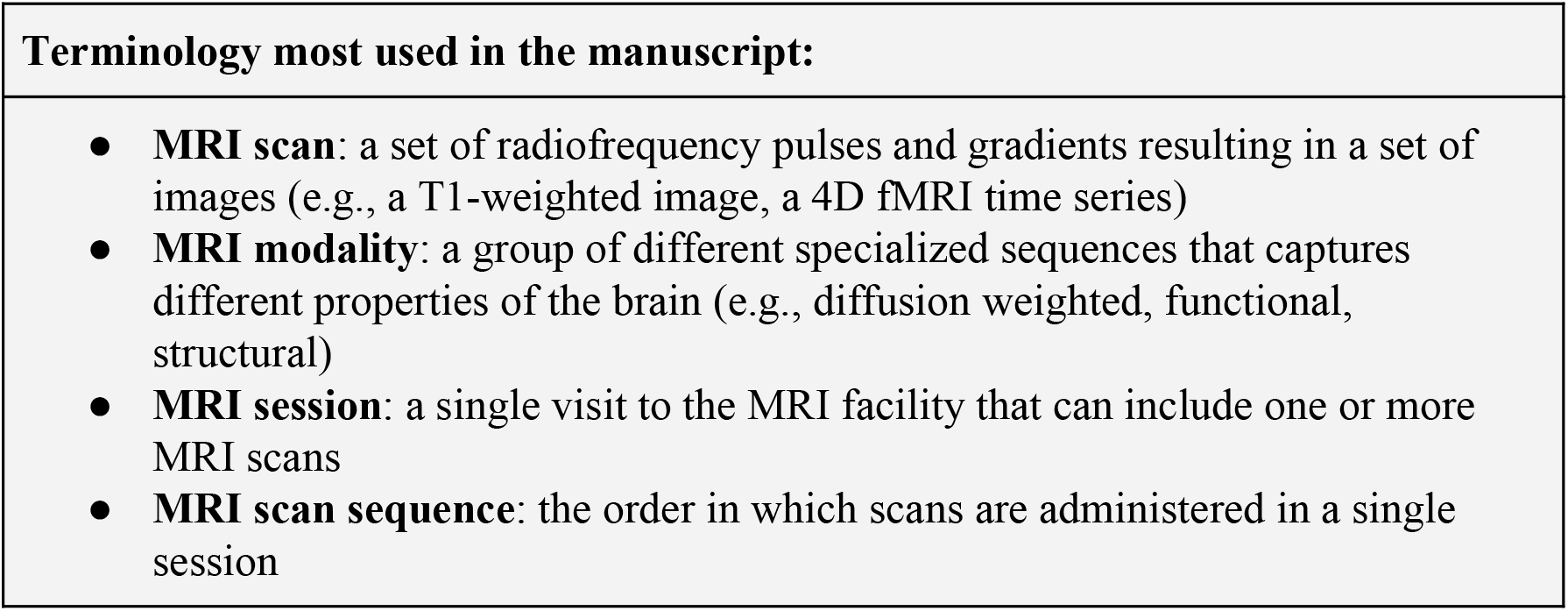
Definitions for selected terms most used in this report.

### 2.4 MRI data quality assurance

For data quality assurance (Q/A) of the T1-weighted and T2-weighted structural scans, images were visually inspected for any motion artifacts or abnormalities, such as blurring, ghosting, or Gibbs ringing artifacts and marked as passing Q/A by either one of two visual reviewers with excellent inter-rater (see Supplementary Methods). Rest and task functional images were also visually inspected for signal dropouts or artifacts and motion was indexed by median framewise displacement (FD) (Jenkinson et al. 2002). Resting state fMRI scans with a median FD≤0.2mm were considered passing Q/A. For task fMRI scans, a cutoff of median FD≤0.4mm was used, given the relative robustness of task-related fMRI designs (Johnstone et al. 2006; Siegel et al. 2014). Diffusion-weighted images were preprocessed with the DTIPrep software package (Oguz et al. 2014). As described in Supplementary Methods they were considered passing Q/A if more than 50% of the gradients collected met our quality criteria.

### 2.5 MoTrak sensor data preprocessing

The recorded MoTrak sensor data were quantized to match the sensor’s measured resolution of 0.77 mm and 0.2°. Similarly, motion introduced when re-centering the sensor following subject motion or positional drift was also removed. For each child, the mean FD was calculated from the motion recorded during the final six minute MRI simulator session and used in statistical modeling analyses. When comparing recorded motion from those who passed the MRI simulator protocol versus those who failed, the last available motion recording was used. If a six minute recording was not available, the longest data section available before that was used.

### 2.6 Statistical analysis

#### Sample characterization

Groups (i.e., ASD vs. ADHD, passing vs. failing MRI simulator training) were characterized and compared in regard to clinical symptom severity and demographics using Mann-Whitney U (Mann and Whitney 1947) or chi-squared tests for continuous and categorical variables, respectively. To correct for multiple comparisons, we used the Benjamini-Hochberg false discovery rate correction (Benjamini and Hochberg 1995) with an alpha of 0.05.

#### Predictive feature selection

We explored the importance of features predicting the ability to successfully complete a MRI multiscan session among a range of child characteristics and performance at the MRI simulator training. Training performance features included mean FD of the final six-minute training step and the number of training sessions needed to pass the MRI simulator training. Child characteristics included age, intelligence quotient (IQ), and severity indexes of ASD and ADHD core symptoms, as well as associated psychopathology symptoms. Given the frequent overlap and co-occurrence of psychiatric symptoms across ASD and ADHD (Reiersen and Todd 2008; Simonoff et al. 2008; Rommelse et al. 2010; Grzadzinski et al. 2011; Leitner 2014; Kern et al. 2015; Joshi et al. 2017), we leveraged a range of parent and clinician based instruments providing continuous measures across groups (see list in Box 2 and Supplementary Methods). To capture the distinct components involved in the symptom and cognitive domains of interest, we selected subscale scores. For children for which one or two of the instruments used to derive the features were missing (n=11 ASD, n=15 ADHD w/o ASD), missing values were imputed. For imputation, we computed the mean value of the missing measure from available data in children matched by both age and diagnosis with those missing the metric to impute. Children with missing data for more than two instruments were excluded from these analyses (n=11).

#### Predicted MRI outcomes

Any given MRI scan completed and passing Q/A was considered successful (passing). Scans not meeting Q/A criteria, incomplete or not attempted were considered failing. Task fMRI scans that were not attempted because the child failed the task practice before the session (n=23, see Supplementary Methods) were considered failing in this context. A post hoc analysis showed that inclusion of data from these scans did not confound results (data not shown).

**Box 2.**
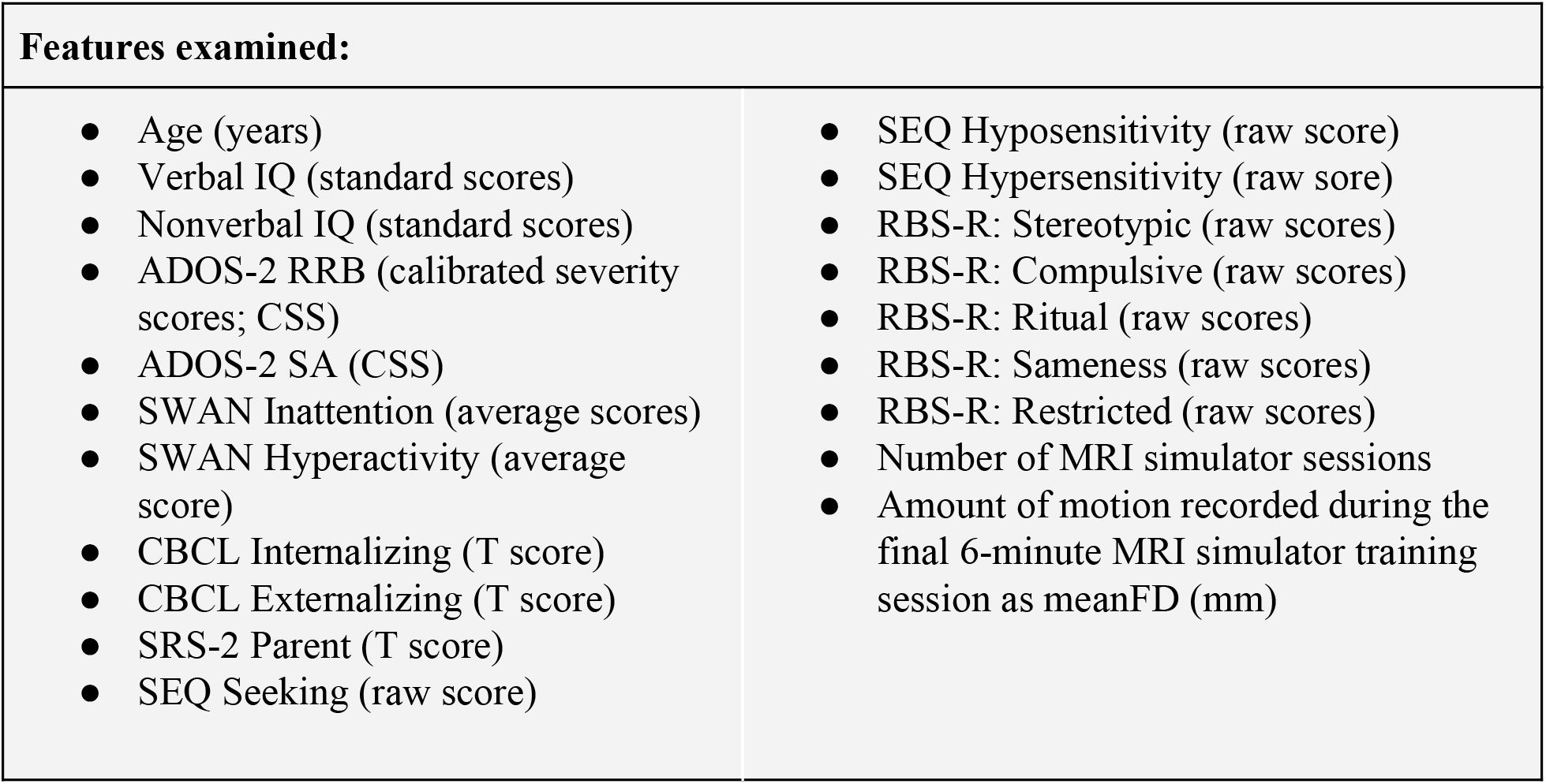
Features examined to predict MRI scan outcomes. Abbreviations: ADHD, attention-deficit/hyperactivity disorder; ADOS-2, Autism Diagnostic Observation Schedule - second edition; ASD, autism spectrum disorder; CSS, calibrated severity scores; CBCL, child behavior checklist; RBS-R, Repetitive Behaviors Scale – Revised; RRB, restricted and repetitive behaviors; SEQ, Sensory Experience Questionnaire; SRS-2, Social Responsiveness Scale-second edition; SWAN, Strengths and Weaknesses of ADHD-symptoms and Normal-behaviors.

#### Random forest (RF) regression

was used to assess which factors influenced the number of scans a child successfully completed in a given MRI session. Predicted values ranged from zero (not successfully completing T1w and subsequent scans) to six (successfully completing all scans), with intermediate values representing completion of T1w plus each of the subsequent scans along the sequence illustrated in Figure 2. For the purposes of these analyses, the two task-fMRI scans shown in Figure 2 were combined such that six scans were assessed (whereby failing reflected failing both scans). The inherent properties of RF, including flexibility regarding input feature types, lack of overfitting, and associated feature importance methods made it an appropriate choice for our question. For this report, we used the scikit-learn random forest implementation (Pedregosa et al. 2011). The RF was trained with default parameters listed in this manuscript’s GitHub repository along with the full code (github.com/aksimhal/mri-simulator-analysis); the number of estimators was set to 300 to increase the performance of the regressor. Results were obtained by training a RF with stratified five-fold cross validation, repeated 100 times.

Feature importance was calculated using the permutation importance method (Breiman 2001). Briefly, we recorded a baseline accuracy score for the trained regressor, permuted the values of each feature, then passed all the test samples back through the RF and recomputed accuracy. The importance of a given feature was indexed by the difference between the baseline and the new accuracy value (Breiman 2001); this is known as the ‘out of bag error’ (OOBE). A feature or a set of features were selected as important when their OOBE had the largest gap from the following feature’s OOBE in the model rank.

#### Naive Bayes classification

To assess what factors contributed to completing a minimal multiscan dataset, we examined the children who completed our first T1w structural scan and following R-fMRI scan with data passing Q/A. Because of the imbalanced nature of the dataset (88% of those who attempted both scans, passed), we used the naive Bayes (NB) classification method, which takes into account posterior probabilities. The NB classification implementation used was from scikit-learn (Pedregosa et al. 2011). The NB model was trained using the 20 features listed in Box 2 to predict whether or not the subject successfully completed both the T-w sequence and the R-fMRI scan. Results were obtained via five-fold cross validation, repeated 1000 times. Feature importances were calculated using the same permutation method adapted from (Breiman 2001) as described above.

## 3. Results

### 3.1 Characteristics of the sample

We examined data from 250 children who attempted to complete at least one MRI simulator training session. As shown in Supplementary Table 3, n=112 (46%) children had a primary diagnosis of DSM-5 ASD (with or without ADHD comorbidity) and n=138 (55%) had a DSM-5 ADHD primary diagnosis, with or without any other comorbidity but no ASD - here referred as ADHD_w/oASD_. Characteristics of the sample in regard to demographics and clinical presentations are in Supplementary Table 1. Briefly, as expected, children with ASD showed significantly more severe ratings of ASD symptoms relative to those with ADHD_w/oASD_. On the other hand, and likely due to the high comorbidity rates of ADHD in ASD (45%), ADHD symptom severity parent ratings did not statistically differ between the two diagnostic groups. Although children with ASD had higher severity ratings of associated psychopathology, as indexed by the parent CBCL T scores, this difference did not reach statistical threshold. Notably, full scale and verbal IQ were significantly lower in ASD, albeit both groups’ averages were in the typical intelligence range.

### 3.2 MRI simulator training outcomes

The flowchart in Figure 3A shows the number of children going from attempting at least one MRI simulator training session to the MRI scan appointment. As shown in the barchart in Figure 3B, 150 children (60%) successfully completed the training in one session. We invited the remaining 100 children to return for further training, 88 children accepted the invitation. Of them, 71 (80%) successfully completed the additional MRI simulator training, most on a second session (see Figure 3B and Table 2) yielding a total of 221 children passing the simulator training (Figure 3A). Among the 29 (12%) who failed to complete the MRI simulator training, six declined to enter the MRI simulator; the others attempted one or multiple steps of the training. All children successfully completing the MRI simulator training were invited to participate in an MRI session and 201 children attempted it. The average time between the successfully completed MRI simulator session and the actual MRI appointment was 15±13 days.

**Figure 3.**
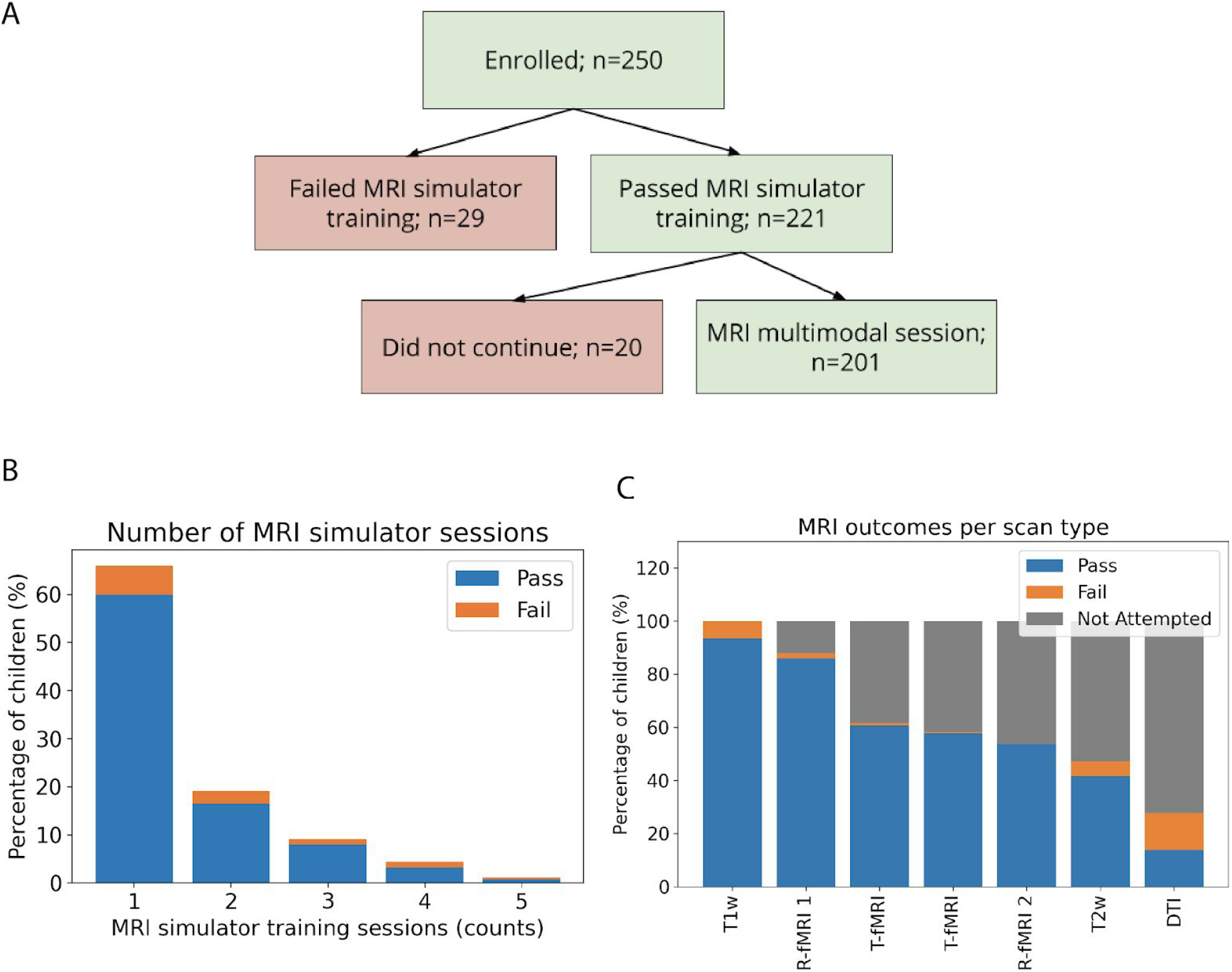
Overview of MRI simulator training and MRI scan outcomes. *A)* Flowchart of participant outcomes. Of the 250 children enrolled in this study, 201 passing the MRI simulator training attempted the MRI multiscan session. *B)* Stacked bars show the number of MRI simulator training sessions needed for children passing the training protocol (blue) vs. those failing it (orange) among the N=250. Most who passed the training (n=150) did so in one session. Of the remaining, n=41 passed training after two training sessions, n=20 after three, n=8 after four, and n=2 after five sessions. Among those who failed the training protocol, n=15 children failed after one, n=7 after two, n=3 after three, n=3 after four, and n=1 after five sessions. *C).* The stacked bars show the percentage of children who attempted each scan with passing or failing Q/A (blue and orange, respectively), the gray stacks represented the percentage of children who did not attempt a given scan along the session. As detailed in the Supplementary Methods, for the task fMRI runs, 23 (11%) children were unable to complete the practice tasks outside the scanner and thus were not administered the task fMRI. Information for seven children regarding task practice was not available and they were counted among those not attempted.

As detailed in Table 2, Mann-Whitney U comparisons of demographics, clinical characteristics and MRI simulator training performances of those passing (n=221) and failing (n=29) the MRI simulator training showed that those who passed were on average older, had a higher verbal and non-verbal IQ, and less severe autistic traits. Notably, ADHD symptom severity, indexed by parent SWAN ratings, did not statistically differ among the two groups. A primary DSM-5 diagnosis of ADHD_w/oASD_ was the most frequent among children passing the MRI simulator training.

### 3.3 MRI scan outcomes

Of the 201 children who passed the MRI simulator training and agreed to attempt the MRI multimodal session, nearly all (n=188, 94%) were able to successfully (i.e., data collection passed Q/A) complete at least the first scan in the sequence (T1w). As shown in Figure 3C, the percentage of children successfully completing additional scans decreased as the scan sequence progressed. Figure 4 shows the motion indices across the fMRI data collected, as well as the number of DTI gradients passing Q/A.

**Figure 4.**
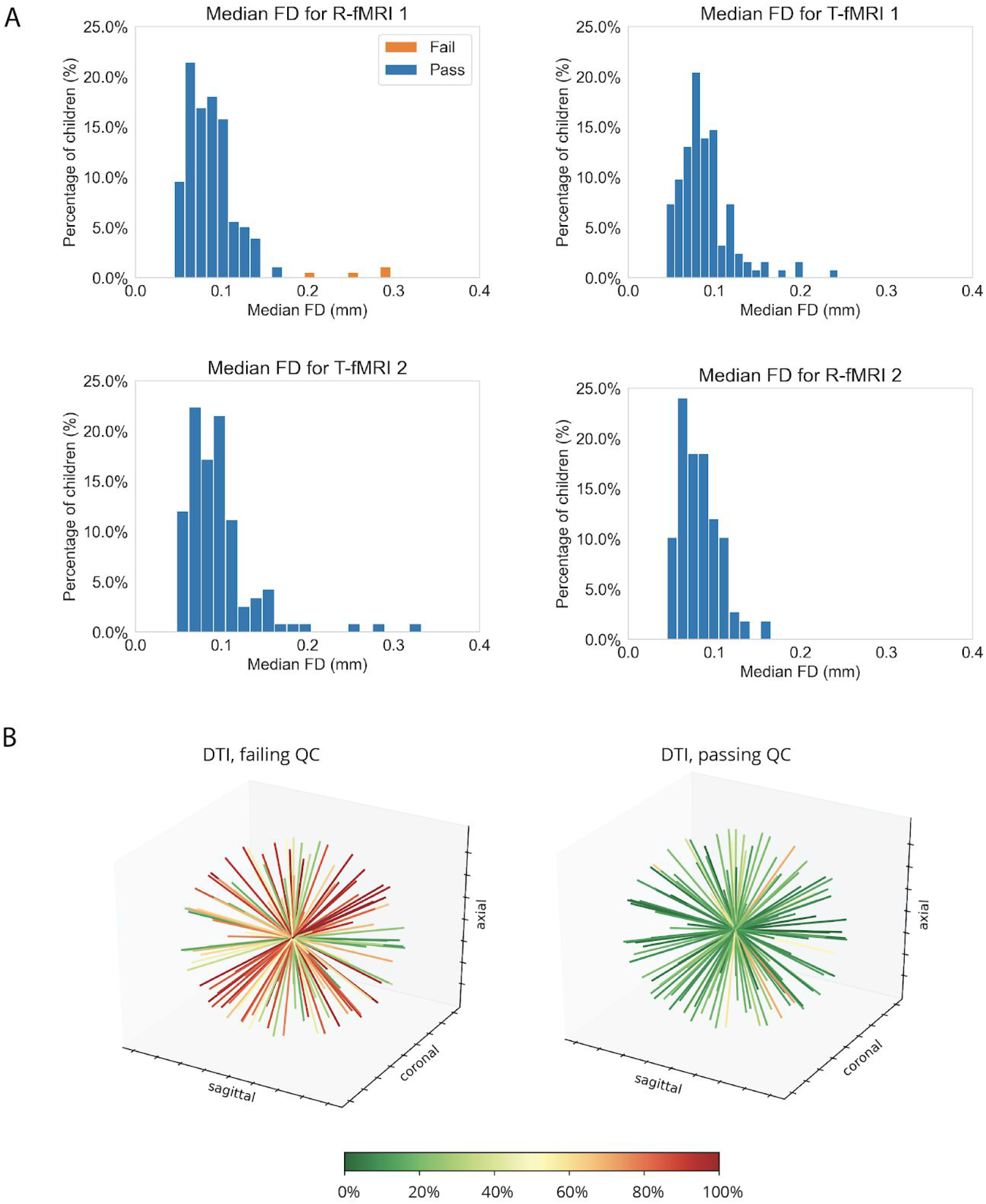
MRI Q/A outcomes. *A)* Histograms of the median frame wise displacement of the subjects who attempted scans. The blue bars represent those who passed and the orange bars represent those who failed. ****B)**** Each plot shows for each gradient direction (represented by a line in 3D), the percentage of participants (represented by the color of the line) with data of sufficient quality as detailed in Supplementary Methods. The plot on the left illustrates the group failing, the one on the right those passing DTI Q/A (n=30 and n=28, respectively).

### 3.4 Predictive features of multiple scans in a MRI session

A random forest (RF) regression was used to examine a combination of clinical, demographic, and mock performance among the 20 features selected (Box 2) that best predicted participants’ degree of successful scan completion (number of completed scans passing). Predicted values ranged from zero (failed T1w and subsequent scans) to six (passed all scans), with any intermediate value representing completion of T1w plus each of the subsequent scans along the sequence described in Figure 2 and in Supplementary Methods.

The mean average error of the trained RF model predicting the number of scans completed was 1.29 scans and the percentage variance explained by the RF model was 17.4%. As shown in Figure 5A, age was the most important feature predicting the number of scans completed in a session, as computed via the permutation importance method described above (i.e., the jump from age to the next feature was 20%). As shown in Figure 5B, children 8.9 ± 1.7 years and older were more likely to successfully complete multiple scans in the MRI session. There was approximately one year difference between those who completed at least the first three scans (M ± SD, 9.3 ± 2 years) versus those who completed the entire set of scans in the sequence (M ± SD, 10.0 ± 3 years). Figure 6 provides the success rate by age for the MRI scan expected along our sequence. The percentage of children below age nine years passing at least the first two scans (i.e., T1w and R-fMRI) was 78% vs. 92% of those aged nine years and older. The success rate of those younger than nine years further decreased with more scans required in the sequence, dropping to 3% for the whole set. A similar pattern was noted for those older than nine years, but with different magnitude (e.g, 20% completed the whole session; see Figure 6). Further, secondary analyses adding diagnostic category labels (ASD and ADHD without ASD) in addition to the other 20 features as predictors, yielded highly similar results with no noticeable improvement in classifier performance (Supplementary Figure 1).

**Figure 5.**
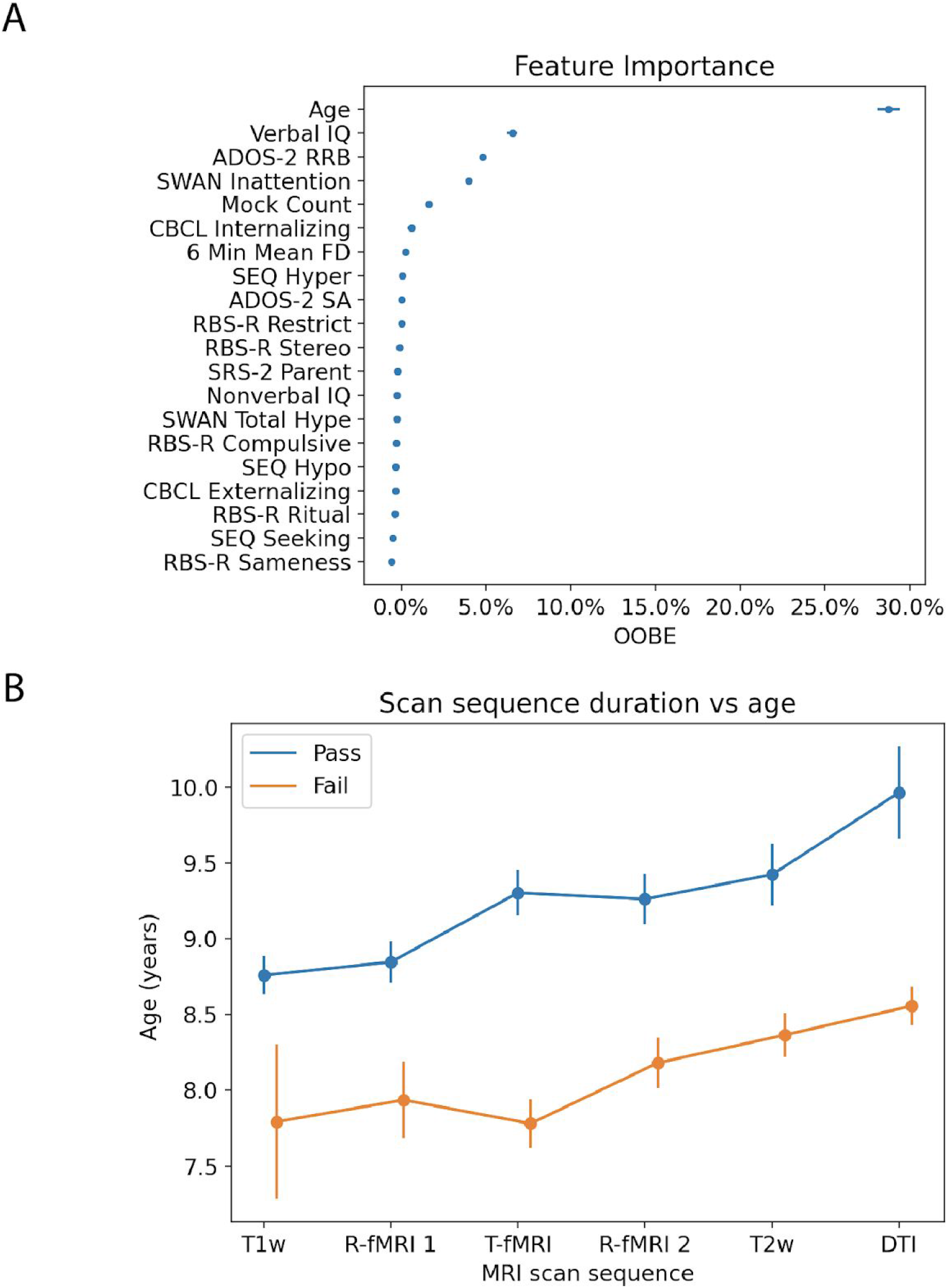
Overview of the random forest regression results. A) Permutation error (feature importance) associated with each of the 20 features examined shown as mean OOBE and standard error across RF-R iterations. B) Group mean and standard errors bars ages for those who passed (blue) and those who failed (orange) along each of the scans in the MRI sequence.

**Figure 6.**
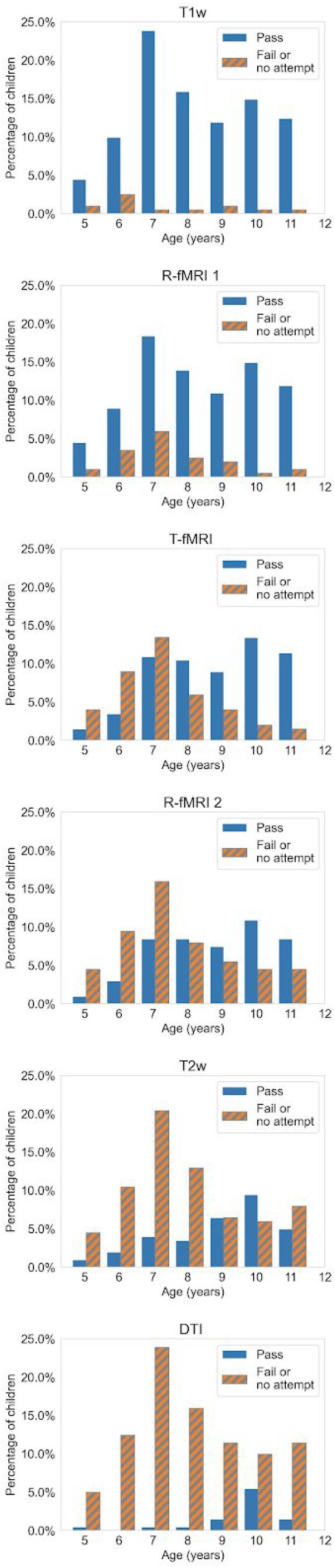
Histograms showing MRI outcome by age. Each histogram shows the percentage of children in each age group who passed or failed Q/A for a given MRI scan in the sequence. The order in which scans were conducted is shown in Figure 2.

Additionally, following age, three features (VIQ, ADOS-2 RRB CSS, and SWAN inattention scores) clustered together before the next OOBE jump, which was 0.02%. As such, we explored the distributions of these variables between children failing and passing each MRI scan as a function of the MRI scan sequence order. As shown in Supplementary Figure 2, children failing scans on average had notably lower VIQ and more severe RRB ADOS-2 scores. SWAN inattention scores on average were slightly higher (more severe) in children failing all scans except T1w and DTI. This pattern of results was consistent even after regressing out age (data not shown). Given that the ADOS-2 RRB scores encompass both “lower- and higher-order” RRB symptoms (Bishop et al. 2013), we explored the pattern of ADOS-2 module 3 items tapping into ‘lower-order’ RRB (i.e., sensorimotor) separately from the item assessing “higher-order” RRB (see Supplementary Methods). As shown in Supplementary Figure 3, while the group passing did not differ in the “higher-order” RRB item score from those failing, notable differences were evident for “lower-order” RRB, even after controlling for age.

### 3.5 Predictors of minimum set of multimodal scan success

To assess which factors influenced the ability to successfully complete a minimum set of multimodal scans (here T1w and R-fMRI scan), a naive Bayes classifier assessed the 20 features listed in Box 2 to predict the labels “pass” vs. “fail.” The resulting classifier had an average accuracy of 74.6%, average recall of 86.3%, and average precision of 84.0%. The true positive rate was 0.729, the true negative rate was 0.017, the false positive rate was 0.138, and the false negative rate was 0.116. The RBS-R Stereotype subscale score was ranked as the most important feature, albeit with an out-of-bag-error (OOBE) score of 0.19%, as shown in Figure 7A. The next most important feature was the 6 minute meanFD measurement from the last MRI simulation training step with an OOBE score of 0.17%, which was followed by VIQ (OOBE score of 0.14%). Remaining features in the rank had smaller OOBE (<0.10%) and thus were not considered to be important. When we examined the average distribution of the RBS-R Stereotype subscale, more severe scores characterized children failing (M±SD, 3.3 ± 3.4) versus those passing (M±SD, 2.6 ± 2.7) the minimum set of scans (T1w + R-fMR1), as shown in Figure 7B. Notably, RBS-R subscale scores indexing “higher-order” RRB (Sameness, Restrictive, Ritual, and Compulsive) were largely associated with negative permutation errors, implying that they did not contribute to the successful completion of these scans. As shown in Supplementary Figure 4, while lower VIQ also predicted passing the minimal set of scan success, here, lower motion at the “mock scan” characterized children passing vs those failing the T1w + R-fMR1 set. Secondary analyses adding diagnostic categories among the 20 continuous variables included as predictors in the naive Bayes classifier, yielded highly similar results to the primary analysis, with no noticeable improvement in classifier performance (Supplementary Figure 5).

**Figure 7.**
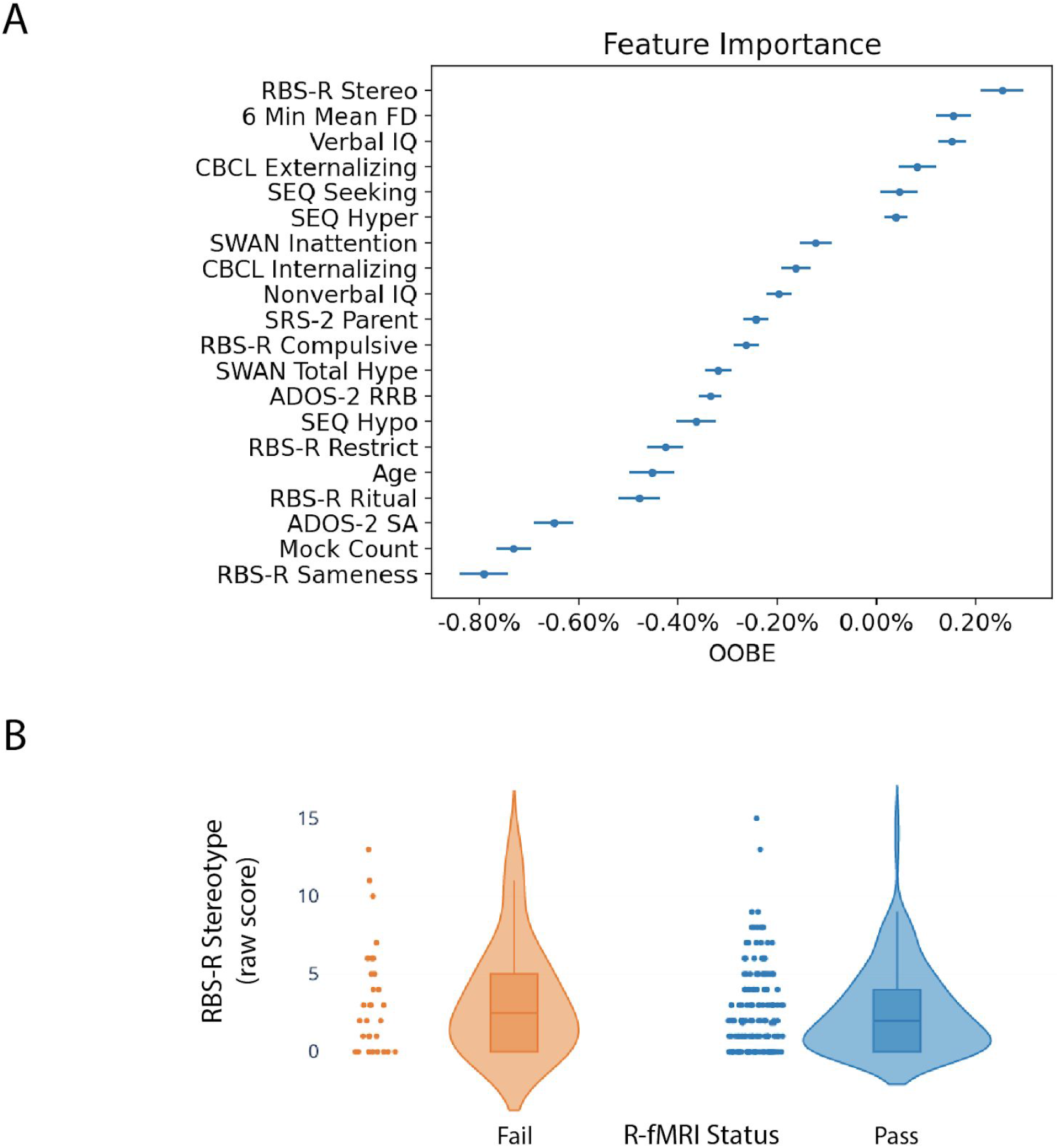
Overview of naive Bayes-based results. A) The top panel shows the permutation error (feature importance) associated with each variable examined. As mean and standard error across the classifier iterations. B) The bottom panel shows the distribution of Repetitive Behaviors Scale-Revised (RBS-R) Stereotype subscale raw scores as violin plots for those who passed (blue) and those who failed (orange) to successfully complete the T1w+R-fMRI. Each dot on the scatter plot indicates a child’s score. The violin plots model the distribution of the scores. The boxplots show the quartile ranges of the data.

## 4. Discussion

We examined factors contributing to the successful collection of multiple scans in a relatively large sample of verbally-fluent school-age children with ASD and/or ADHD. Although a substantial number (88%) of participants successfully completed at least the first two MRI scans in the sequence (T1w + R-fMRI), the success rate decreased with the number of scans collected in the MRI session. This was inversely related to child age, regardless of diagnosis. Beyond age, sensorimotor atypicalities indexed by lower-order RRB ratings, verbal cognitive skills, and pre-scan MRI simulator training performance predicted successful imaging, again, regardless of diagnosis. Collectively, these findings can inform decision-making regarding scan protocols in neurodevelopmental conditions, based on expectations regarding quality data yield.

### 4.1 Age impacts the number of scans completed

Our findings that age is the most important predictive feature of MRI scans success in a session are consistent with prior studies in typically and atypically developing children (Yerys et al. 2009; Rajagopal et al. 2014), which related higher scan failure rates to younger ages. Those prior studies examined the impact of age on one or up to three scans in an MRI session. Here, we extended the scope by examining multiple structural and functional scans in a session. We found that, across all children, the more scans required to be collected in the session, the higher the number of scan failures. Notably, the failure rate was higher in children younger than nine years. For example, the success rate gradually dropped from 95% for the first scan to 20% for the last and seventh scan required in the sequence in those older than nine years. In contrast, the success rate decreased from 78% for the first to 3% for the last scan in children younger than nine years.

These results suggest that brain imaging research focusing on school-aged children with ASD and/or ADHD needs to account for differential age-related attrition, particularly as the number of scans intended for a session increases. This may require over-recruiting younger individuals to adjust for greater attrition rate, and/or adjusting the age range targeted, or limiting the number of MRI sequences obtained in a single session, or considering multiple MRI sessions in the design. We do not point to a single solution as this may vary as a function of context, resources, and goals in research or clinical settings where multiple scans and/or multimodal imaging are increasingly required. An important implication of the present work for the clinical setting is its demonstration of successful awake imaging for children with ASD and/ or ADHD aged nine and over. Consistent with prior reports ((Rosenberg et al. 1997; Nordahl et al. 2008; Barnea-Goraly et al. 2014) our findings support that investing in MRI simulator training may be an alternative to using sedation for clinical imaging - even in children with neurodevelopmental conditions and extends this notion to multiple scan sessions.

### 4.2 Sensorimotor atypicalities impact scan performance

Notably, while age strongly predicted the extent to which children completed multiple scans in an MRI session, other clinical factors were most important for completing a minimal set of scans (T1w + R-fMRI). These included the severity of restricted and repetitive behaviors (RRB) indexed by parent ratings at RBS-R Stereotype subscale, which measure the severity of sensory and motor RRB (Bodfish et al. 2000; Lam, Bodfish, and Piven 2008). RRB is an umbrella term that refers to a range of symptoms that can be summarized as ‘higher-order’ behaviors, such as insistence on sameness, and ‘lower-order’ behaviors, such as hyper- or hypo-sensitivity to sensory stimuli, as well as motor stereotypies (Bishop et al. 2013). Our RBS-R results revealing a predictive role of the Stereotype scale but not for the other subscales, suggest that ‘lower-order’ RRB — i.e, sensorimotor atypicalities - have a greater role than ‘higher-order’ RRB. Interestingly, after age, ‘lower-order’ RRB, indexed by clinician-based ADOS-2 item scores of sensorimotor atypicalities, contributed to the top four features predicting the number of scans completed in a session. The convergence across analyses onto ‘lower-order’ RRB symptoms, further underscores the role of sensory and motor control processes across children. Although RRBs are considered core ASD symptoms, they have been increasingly reported in a subsample of children with ADHD (Martin et al. 2014). This evidence, combined with our findings that MRI completion was not particularly driven by primary diagnosis, underscores the transdiagnostic impact of sensorimotor symptoms in MRI outcome prediction.

Given that the RRB ratings found to be predictive included both sensory and motor control atypicalities, their relative contribution remains somewhat unclear. On one hand, it is plausible that the sensory experience of being in the MRI scanner (sounds, touch, and visual stimuli) impact children with greater sensory atypicalities, even after comprehensive MRI simulation training. If so, studies examining the role of specific sensory modalities may be relevant to design habituation techniques for school-age children with ASD and/or ADHD. On the other hand, the observation that parent ratings focusing on sensory processes alone (i.e., SEQ) were not predictive features, suggests that the combination of both atypical motor and sensory processes may be more relevant. With this in mind, and given the transdiagnostic nature of our findings, other measures of sensorimotor atypicalities may be explored in future studies of neurodevelopmental conditions.

Looking forward, an increased focus on neurological soft signs (NSS) may provide additional insight on the predictive role of sensorimotor processes. NSS encompass clinically detectable poor motor coordination, sensory perceptual difficulties, and involuntary movements. An emerging literature underscores the role of neurological soft signs as markers of brain immaturity across multiple psychopathologies (Martins et al. 2008). Indeed, NSS are frequent in both children with ASD, those with ADHD, and other neurodevelopmental conditions (Patankar et al. 2012; Manouilenko et al. 2013). Our study did not include a measure of NSS, e.g., Physical And Neurological Examination of Soft Signs (Camp et al. 1977) and to date, the relation between such a measure and RRB metrics is unknown. Thus, future research examining their unique or relative contribution predicting MRI success is needed.

### 4.3 Predictive value of motion during MRI simulator training for scan success

Interindividual variability in meanFD of motion tracking during the last 6 minute step of our MRI simulator training protocol contributed to the prediction of success rate for the minimum first set of multimodal scans (T1w + R-fMRI). This finding underscores the utility of motion tracking during MRI simulator training. We note that the motion sensor hardware available in our study was not designed to capture subtle motion, thus future studies may benefit from improved sensor hardware. Beyond its utility in providing real-time objective feedback associated with a motion event, results from motion tracking may guide decisions on further training prior to a MRI scan visit. Although the number of MRI simulator training sessions did not robustly predict MRI scan success, we note that by giving children the option to undergo additional MRI simulator training, we were able to increase the yield of children successfully completing training from 60% in one session to 88% with multiple training sessions. We note that a recent study from a small sample of children with ASD showed a greater rate of MRI success in children completing a more systematic training versus those completing a less structured one (Horien et al. 2020). Together with our results these observations suggest that greater exposure and habituation is helpful and that this should be accounted for when designing neuroimaging studies of neurodevelopmental conditions.

### 4.4 Limitations and considerations for future studies

The results of this study need to be interpreted in light of several limitations. First, we did not include a training comparison condition-group to directly assess the efficacy of the MRI simulator protocol. Given the large literature supporting the role of MRI simulator training for MRI completion across ages (Greene et al. 2018; Barnea-Goraly et al. 2014; Nordahl et al. 2016; De Bie et al. 2010; Cox et al. 2017; Carter et al. 2010), this was not considered practical, nor cost-effective for our larger ongoing neuroimaging study. Second, we did not examine factors more directly related to the MRI session that may have played a potential role in MRI collection success in our protocol (e.g., number of scans repeated, feedback provided). This was motivated by the explicit goal to identify factors that may guide decisions before scheduling a scan session. A range of approaches aimed to improve data collection during scans continue to emerge (Vanderwal et al. 2015; Dosenbach et al. 2017; Ai et al. 2020; Power et al. 2019). Each has strengths and weaknesses and no one is applicable to a range of different scans and populations to date, unlike MRI simulator training. For example, FIRMM has been shown to be useful for functional MRI data collection (Dosenbach et al. 2017) however, its efficacy with structural and diffusion MRI data is yet to be established. Scans including prospective motion correction have been shown promising for structural scans (e.g., T1-weighted) (Ai et al. 2020). CaseForge has been shown to be effective at reducing head motion in small samples of children and two adults completing R-fMRI (Power et al. 2019; Lynch et al. 2021), but not in another sample of adults completing task-fMRI (Jolly, Sadhukha, and Chang 2020, 2021). The role of CaseForge in limiting motion for longer multimodal scan sessions including different structural scans is also unknown. Passive movie viewing is largely used for structural MRI data collection and is increasingly becoming another promising method for functional scan success in challenging populations (Vanderwal et al. 2015; Eickhoff, Milham, and Vanderwal 2020). Third, because our MRI sequence order was quasi-fixed, we could not assess whether a specific MRI modality along the sequence may be associated with greater success rate. Thus, it remains unclear if the lowest success rate for DTI scans was solely the result of being the last scan in our session or other factors associated with this sequence (louder noise, higher table motion, longest duration) are involved. Similarly, our third set of scans included two short task fMRI blocks involving a relatively simple two-choice matching task selected from the HCP sequence (Hariri et al. 2002; Barch et al. 2013). For these task-fMRI scans, the rate of success among children below nine years of age was lower than chance (50%) suggesting that including task-fMRI in a multiscan neuroimaging session may not be feasible for younger children with ASD and/or ADHD. Fourth, in the statistical analysis, the imbalance of children who passed and failed at each step of the sequence may have limited the predictive power of the models. We mitigated these effects by using five-fold cross validation when evaluating performance and by running the model multiple times to determine aggregate performance. Finally, as the present study focused on verbally fluent children with ASD and/or ADHD with intelligence ranging from borderline to high, future studies should assess the unique challenges presented by younger non-verbal or minimally-verbal children in both the MRI simulation training and MRI scan. This is particularly notable, given that verbal IQ was among the top predictive factors across analyses after age and sensorimotor atypicalities. Further, unique challenges also arise in multimodal imaging of infants, toddlers and preschoolers younger than five. Promising approaches with natural sleep scans or awake scanning (Ellis et al. 2020) are emerging.

## 4.5 Conclusions

In summary, this report provides a first comprehensive assessment of factors predicting success rate in multimodal imaging in verbally fluent school-age children with ASD and/or ADHD following a systematic MRI simulator training. The methods described and the demonstration of the role of age and other clinical features may provide useful insights in designing MRI pediatric protocols in school-age children with ASD and/or ADHD.

## Acknowledgements

The authors are very thankful to all children and their parents/caregivers who generously volunteered their time. The authors also wish to thank the research staff of the Autism Center at the Child Mind Institute and of the Autism Research and Clinical Program of the Child Study Center at NYU Langone Health, who contributed to aspects of recruitment, assessments, and data collection and management as well as Drs Maarten Mennes and Amy Roy for their inputs on earlier version of the MRI simulator training. The authors are also grateful to Dr. Pablo Velasco and the staff at the NYU Center for Brain imaging for their technical and administrative support in MRI data collection.

