## Supplementary information for "Predicting multimodal MRI outcomes in children with neurodevelopmental conditions following MRI simulator training"

**Supplemental information**

***Contents:***


- Supplementary Methods
    1. Sample recruitment and characterization
    2. MRI scan protocol
    3. MRI data quality assurance (Q/A)

- Supplementary Figures
    1. Features importance after adding diagnosis to the random forest regression model
    2. Secondary predictive features associated with scan success in the first MRI session
    3. Age-Residualized ADOS-2 lower- and higher-order RRB item scores
    4. Secondary predictive features associated with the minimal set of scan success
    5. Feature importance after adding diagnosis to the naive bayes classification model

- Supplementary Tables
    1. Characteristics of the children by enrolling sites
    2. MRI scan parameters
    3. Study sample characteristics

**Supplementary methods**

**1- Sample recruitment and characterization**

Participants in this study were recruited through clinical and community referrals from the NYU Grossman School of Medicine and from the Child Mind Institute clinical practices, school psychologists, web postings targeted to parent and advocacy groups, emails among participants in Simons Foundation Powering Autism Research (SPARK), direct mail advertising, and word-of-mouth. As detailed elsewhere (Guttentag et al. 2021), study inclusion required a clinician-based DSM-5 primary diagnosis of ASD (with any comorbidity) or ADHD (with any comrondoty but ASD- i.e., ADHD without ASD; $ADHD_{w/oASD}$), total IQ>65 and fluent English speaking. Exclusion criteria for all participants were genetic syndromes known to be associated with autism, other medical illnesses requiring chronic treatment, and use of antipsychotics within the past six months. No individuals were excluded based on sex, parent-reported race, ethnicity, nor socioeconomic status, indexed by the Hollingstead system (Hollingshead 1975). Children treated with stimulants were asked to withhold them for at least 24 hours prior to testing, MRI scan simulation training, and MRI sessions.

Clinical diagnoses were based on DSM-5 criteria for either one of two primary diagnostic categories: ASD, with or without comorbidities, including ADHD; or ADHD, with or without any comorbidities, excluding ASD - i.e., $ADHD_{w/oASD}$. Diagnoses were supported by parent interviews, direct child assessments, review of available parent questionnaires, and any available prior records. At least two evaluators were involved in the diagnostic evaluation, including licensed psychologists and/or supervised post- and pre-doctoral fellows. One evaluator completed the diagnostic parent interviews that included the Schedule for Affective Disorders and Schizophrenia for School-Age Children-Present and Lifetime Version (KSADS-PL) (Kaufman et al. 1997), and the Autism Screening Interview - school age (Bishop et al. 2017), administered in-person. Another clinician, who was “blind” to prior diagnoses and records, administered and scored the Autism Diagnostic Observation Schedule, second edition (ADOS-2; (Lord 2012), and the Differential Ability Scales-2nd Edition (Elliott 2007). Following these assessments, both the child and parent evaluators discussed their diagnostic impressions and reviewed all available clinical information to reach a consensus on best-estimate clinical diagnosis that was confirmed at a multidisciplinary (i.e, child psychiatrist, child psychologist, social worker) case conference. Additional parent-based measures were collected to further characterize the sample in terms of the severity of core ADHD and/or ASD symptoms, as well as of other comorbid psychopathology to assess predicting features of scan success. To better capture the different components involved in the symptom and cognitive domains of interest, we selected *a priori* valid subscales instead of summary scores. These additional parent-based measures included:

- **Strengths and Weaknesses of ADHD-symptoms and Normal-behavior** (SWAN) rating scale (J. Swanson et al. 2006; J. M. Swanson et al. 2012). The SWAN is a parent/caregiver 18 item questionnaire designed to assess hyperactivity, impulsivity, and inattention on a 4-point Likert scale ranging from -3 to 3 in school age children. It yields an average (and raw total) score for three scales: ADHD, hyperactivity/impulsivity, and inattention index. For the present study prediction analyses, we used the average subscores of the inattention and hyperactivity indices.
- **Child Behavior Checklist** (CBCL) is a widely used parent questionnaire consisting of 113 items. It rates problem behaviors associated with psychopathology on a 3-point Likert scale (Achenbach and Ruffle 2000). It provides T scores on eight syndrome domains and 6 DSM categories. The syndrome domains (*anxious/depressed, withdrawn/depressed, somatic complaints, social problems, thought problems, attention problems, rule-breaking behavior, and aggressive behavior)* are summarized in a total problem summary T-score. Internalizing and externalizing problems subscale summary T-scores are also provided: one combining *anxious/depressed*, *withdrawn-depressed*, and *somatic complaints* scores, the other *rule-breaking* and *aggressive*

*behavior* scores. For the present prediction analyses, we used the T-scores of the internalizing and externalizing subscales.

**Social Responsiveness Scale-second edition (SRS-2)**. The SRS-2 includes 65-item on a 4 point Likert-scale that assesses current (past 6 months) ASD related symptoms, with 53 items focusing on social communication and 12 items on restrictive, repetitive behaviors (RRBs). The resulting total score is converted in total T-scores for boys and girls, separately (Constantino and Gruber 2012). For the present prediction analyses, we used T scores as recommended, given that subscale scores are not sufficiently independent (Constantino 2011)

**Sensory Experience Questionnaire (SEQ;** (Baranek et al. 2006; Little et al. 2011; Ausderau and Baranek 2013)). The SEQ is a parent/caregiver 21-item questionnaire where responses are marked on a 5-point Likert scale. It measures sensory patterns of hyper- and hyper-responsiveness as well seeking behavior in social and non social context provides a total and subscale scores. Here we used the summary scores for the hyposensitivity, hypersensitivity, and seeking.

- **Repetitive Behaviors Scale - Revised (RBS-R).** The RBS-R is a parent/caregiver questionnaire that measures repetitive behaviors/interests in children with ASD. It includes 44 items, scored on a 4-point Likert scale (Lam 2004). For the feature importance analysis, we used the raw subscores on the mostly used RBS-R 6-factor structure (Lam and Aman 2007). All factors were included except for self injurious as the range of scores in this case was very limited due to the infrequency of these symptoms in our sample. As a result the RBS-R subscale s*tereotype compulsive, ritual, sameness, and restricted* were included in the analyses.

**2 - MRI scan protocol**

The MRI scan protocol with the order of scan administration during the first scan session  is described in supplementary Table 1 and in Figure 1 of the main text. At the MRI scan visit, following the review of a social story entitled “Getting brain pictures with an fMRI scan" and a metal screening, children were invited in the MRI room. In the MRI room children were first equipped with a respiration belt from the Biopac Systems Inc (“Data Acquisition and Analysis System - MP160 System - Windows” 2020) and earplugs to reduce in-scanner noise. Then, they laid down on the table in a comfortable supine position. Their head rested on the bottom half of the head coil on a vacuum pillow; padding around the sides of the head helped to restrict head motion. The top half of a 32-channel Siemens head coil was then positioned and a mirror mounted on it allowed to view stimuli back-projected to a screen at the rear of the MRI bore. A cartoon video of the child’s choice was administered during all structural and diffusion scans (T1-weighted, T2-weighted, DTI); a white cross centered on a black screen was shown for the resting state fMRI scans.  Finally, the Hariri task (Hariri et al. 2002) was administered in two blocks consistent with the HCP protocol (Barch et al. 2013).  To enable completion of the face emotion recognition task, button boxes were provided. A pulse oximeter was also used and most often positioned on one  the halluces (big toe) of the child. Eye tracking data was collected during the Task-fMRI runs, whenever possible using an eye tracker camera, EyeLink 1000 (SR Research, Ontario, Canada).

To ensure all children were able to understand the fMRI task instructions, at least one short practice of 27 seconds including six trials of the Hariri task was administered. Two practice runs were administered for those completing the first one with accuracy <67%. Only children able to correctly respond to at least 67% of the trials were administered the task in the MRI session. During the course of the study, to maximize the number of children able to complete the task practice, a longer practice version of 54 seconds was administered adding six more faces selected from the NimsTim battery (Tottenham et al. 2009). Overall, out of the 201 children undergoing the first MRI session, 23 children failed the task practice and therefore did not attempt the task fMRI; task practice information was missing for seven children for whom task-fMRI was not collected.

As described in the main text, the scan runs were administered in a quasi-fixed order, always starting with T1 and finishing with DTI (Figure 2, Supplementary Table 1). Specifically, when children moved or asked for a brea, runs were stopped and then repeated as needed and feasible following encouragement and feedback. The administration of the next scan in the sequence depended on the success of the prior one. During data collection, two MRI operators monitored movement directly,  one via the operator room window, the other via the eye tracker camera positioned at the back of the MRI bore providing a direct sight of the child's eye. During functional scans (rest and task), movement across the six directions was quantitatively monitored in real time using an in-house built MATLAB code. The MRI operator stopped the fMRI runs when motion exceeded a pre-specified threshold (rest = 1mm; task = 2mm) in the first few minutes of the scan, and gave feedback and encouragement to repeat the scan, as feasible.

**3 MRI Data Quality Assurance (Q/A)**

To ensure quality of structural scans (T1-weighted and T2-weighted), at least one of two quality raters (JF and PS) visually inspected all  images for any artifacts (e.g., ringing artifacts, ghosting effects, blurred regions, well defined white and gray matter).

Anatomical quality was assured by visual inspection on a fail vs pass rating. To evaluate concordance among the two raters, 84 (42 T1-weighted and 42 T2-weighted; ~28% of the images) a  intraclass correlation coefficient, ICC, (Shrout et al. 1979) and their 95% confidence intervals were calculated using the psych toolbox version 2.0.9 for R based on a single fixed ratings, 2-way mixed-effects model. The resulting $ICC_{(3,1)}$ was 0.96 (Confidence interval: 0.94- 0.97) indicating excellent inter-rater reliability. For rest and task functional images, following visual inspection for signal dropouts or artifacts, motion indexed framewise displacement (FD) (Jenkinson et al. 2002) was computed. Resting state fMRI scans with a median FD≤0.2mm were considered passing Q/A. For task fMRI scans, given the relative greater robustness of task-related fMRI design to motion (Johnstone et al. 2006; Siegel et al. 2014), a median FD≤0.4mm was used to consider for passing Q/A. Regarding the DTI data Q/A, several quantitative steps were taken on a gradient-wise level.  DTI scans with at least 69 gradients passing our quality assurance (out  of the 137 collected, (50%) were deemed passing. Each step is described below

1. FSL Motion Outliers: Gradients with a framewise displacement >3 mm rejected.
2. DTIPrep Image checking: Participants with incomplete scans (prematurely ended and/or otherwise containing less than 137 total gradients) were identified by DTIPrep (Oguz et al. 2014) and all gradients for such participants rejected.
3. DTIPrep Slice-wise Deviation: This step checked gradients for intensity and brightness artifacts. As recommended by DTIPrep and prior studies (Jovicich et al. 2014; Wang et al. 2017), gradients with an average intensity >3.5 standard deviations above the mean were rejected.

4. DTIPrep Interlace-wise Deviation: This step checked for Venetian blind artifacts and assessed within-volume motion, rejecting gradients with an intensity >2.5 standard deviations above the mean intensity of the volume. Gradients were also rejected with a rotation of 0.5 degrees or a translation deviation of 1.75 mm beyond the average position of the volume. These thresholds reflect those recommended by DTIPrep - as default settings (intensity, rotation) or as a parameter-based calculation (translation, average of voxel measurements).

Constantino, John N. 2011. “The Quantitative Nature of Autistic Social Impairment.” *Pediatric Research* 69 (5 Pt 2): 55R – 62R.

Constantino, John N., and Christian P. Gruber. 2012. *Social Responsiveness Scale: SRS-2*. Western Psychological Services Torrance, CA.

“Data Acquisition and Analysis System - MP160 System - Windows.” 2020. 2020. https://www.biopac.com/product/mp150-data-acquisition-systems/.

Elliott, C. D. 2007. “Differential Ability Scales, 2nd Edition.” *Harcourt Assessment*.

Guttentag, Sara, Somer Bishop, Rebecca Doggett, Rebecca Shalev, Megan Kaplan, Margaret Dyson, Morgan Cohen, Catherine Lord, and Adriana Di Martino. 2021. “The Utility of Parent-Report Screening Tools in Differentiating Autism vs. ADHD in School-Age Children.” *PsyArXiv*. https://doi.org/10.31234/osf.io/9pu7t.

Hariri, Ahmad R., Alessandro Tessitore, Venkata S. Mattay, Francesco Fera, and Daniel R. Weinberger. 2002. “The Amygdala Response to Emotional Stimuli: A Comparison of Faces and Scenes.” *NeuroImage* 17 (1): 317–23.

Hollingshead, August B. 1975. “Four Factor Index of Social Status. New Haven.” *CT: Yale University*.

Jenkinson, M., P. Bannister, M. Brady, and S. Smith. 2002. “Improved Optimization for the Robust and Accurate Linear Registration and Motion Correction of Brain Images.” *NeuroImage* 17 (2): 825–41.

Johnstone, Tom, Kathleen S. Ores Walsh, Larry L. Greischar, Andrew L. Alexander, Andrew S. Fox, Richard J. Davidson, and Terrence R. Oakes. 2006. “Motion Correction and the Use of Motion Covariates in Multiple-Subject fMRI Analysis.” *Human Brain Mapping* 27 (10): 779–88.

Jovicich, Jorge, Moira Marizzoni, Beatriz Bosch, David Bartrés-Faz, Jennifer Arnold, Jens Benninghoff, Jens Wiltfang, et al. 2014. “Multisite Longitudinal Reliability of Tract-Based Spatial Statistics in Diffusion Tensor Imaging of Healthy Elderly Subjects.” *NeuroImage*. https://doi.org/10.1016/j.neuroimage.2014.06.075.

Kaufman, Joan, Boris Birmaher, David Brent, Uma Rao, Cynthia Flynn, Paula Moreci, Douglas Williamson, and Neal Ryan. 1997. "Schedule for Affective Disorders and Schizophrenia for School-Age Children-Present and Lifetime Version (K-SADS-PL): Initial Reliability and Validity Data." *Journal of the American Academy of Child & Adolescent Psychiatry*. https://doi.org/10.1097/00004583-199707000-00021.

Lam, Kristen S. L. 2004. "The Repetitive Behavior Scale-Revised: Independent Validation and the Effects of Subject Variables." The Ohio State University. http://rave.ohiolink.edu/etdc/view?acc_num=osu1085670074.

Lam, Kristen S. L., and Michael G. Aman. 2007. "The Repetitive Behavior Scale-Revised: Independent Validation in Individuals with Autism Spectrum Disorders." *Journal of Autism and Developmental Disorders* 37 (5): 855–66.

Little, Lauren M., Ashley C. Freuler, Marisa B. Houser, Lisa Guckian, Kristin Carbine, Fabian J. David, and Grace T. Baranek. 2011. "Psychometric Validation of the Sensory Experiences Questionnaire." *The American Journal of Occupational Therapy: Official Publication of the American Occupational Therapy Association* 65 (2): 207–10.

Lord, Catherine. 2012. *Autism Diagnostic Observation Schedule, Second Edition (ADOS-2)*. WPS.

Oguz, Ipek, Mahshid Farzinfar, Joy Matsui, Francois Budin, Zhexing Liu, Guido Gerig, Hans J. Johnson, and Martin Styner. 2014. "DTIPrep: Quality Control of Diffusion-Weighted Images." *Frontiers in Neuroinformatics* 8 (January): 4.

Siegel, J. S., J. D. Power, J. W. Dubis, and A. C. Vogel. 2014. "Statistical Improvements in Functional Magnetic Resonance Imaging Analyses Produced by Censoring High‐motion Data Points." *Human Brain Mapping*. https://onlinelibrary.wiley.com/doi/abs/10.1002/hbm.22307.

Swanson, James M., Sabrina Schuck, Miranda Mann Porter, Caryn Carlson, Catharina A. Hartman, Joseph A. Sergeant, Walter Clevenger, et al. 2012. "Categorical and Dimensional Definitions and Evaluations of Symptoms of ADHD: History of the SNAP and the SWAN Rating Scales." *The International Journal of Educational and Psychological Assessment* 10 (1): 51–70.

Swanson, J., S. Schuck, M. Mann, C. Carlson, K. Hartman, J. Sergeant, and R. McCleary. 2006. "Categorical and Dimensional Definitions and Evaluations of Symptoms of ADHD: The SNAP and SWAN Rating Scales." *University of California, Irvine*.

Tottenham, Nim, James W. Tanaka, Andrew C. Leon, Thomas McCarry, Marcella Nurse, Todd A. Hare, David J. Marcus, Alissa Westerlund, B. J. Casey, and Charles Nelson. 2009. "The NimStim Set of Facial Expressions: Judgments from Untrained Research Participants." *Psychiatry Research* 168 (3): 242–49.

Wang, Kun, Zhi Chen, Fan Zhang, Qingxin Song, Canglong Hou, Yixing Tang, Jun Wang, et al. 2017. "Evaluation of DTI Parameter Ratios and Diffusion Tensor Tractography Grading in the Diagnosis and Prognosis Prediction of Cervical Spondylotic Myelopathy." *Spine* 42 (4): E202–10.

**Supplementary Figure 1**

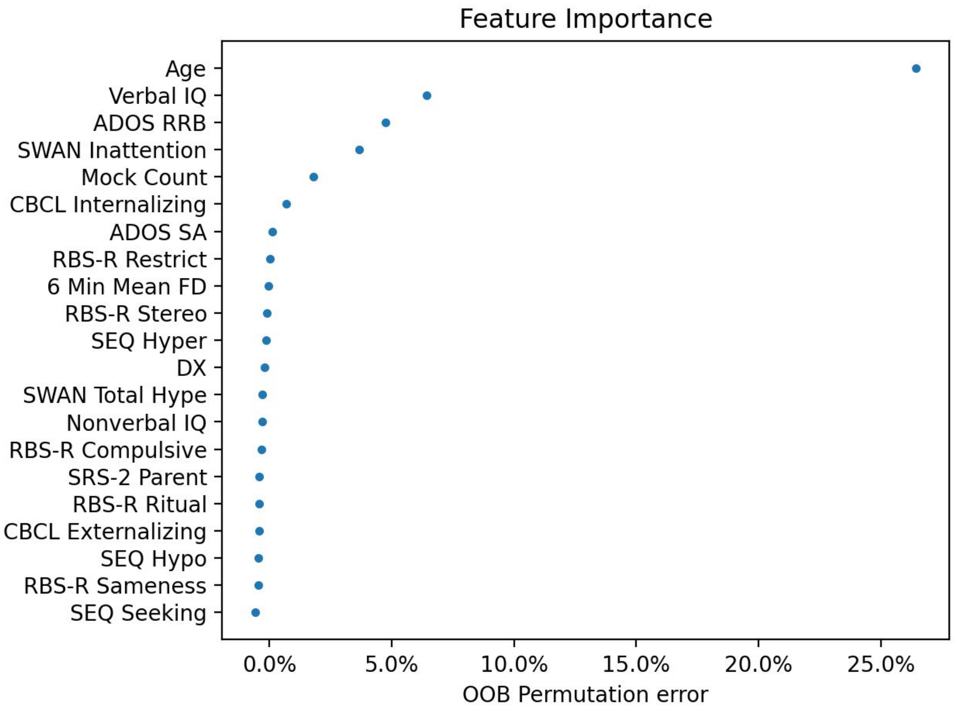


**Supplementary Figure 1. Feature importance after adding diagnosis (DX) to the naive bayes classification model.** Feature importances with the addition of diagnostic label ($ADHD_{w/oASD}$ vs ASD) to the random forest regression model. After adding diagnosis as a feature, the model's virtually did not change (mean average error 1.31 scans, model's variance explained remained around 15.7%).

***Supplemental Figure 2:***

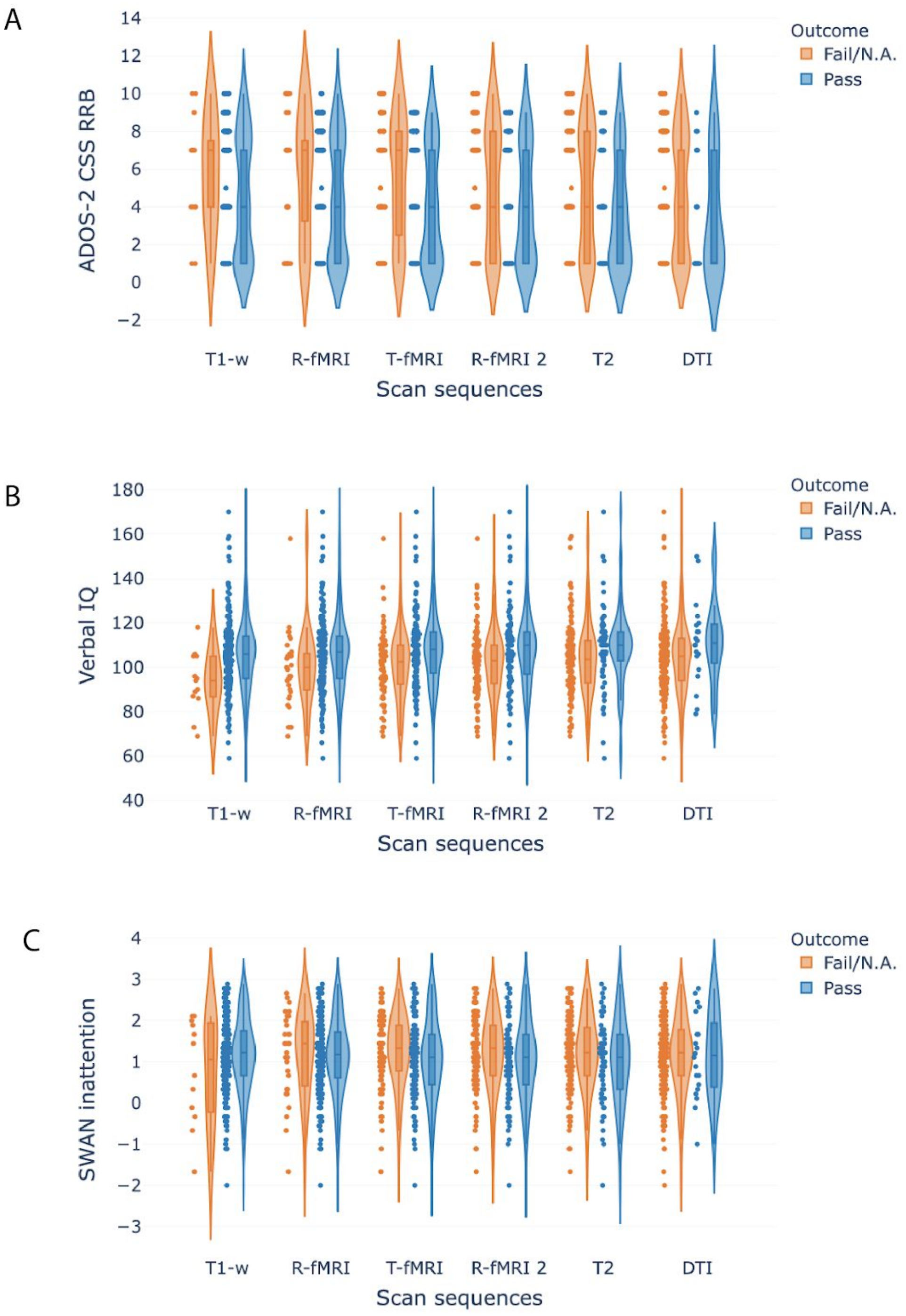


**Supplementary Figure 2: Secondary predictive features associated with scan success in the first MRI session.** Scatter, violin, and box plots show the group distribution of clinical measures for each MRI scan. A) ADOS-2 RRB subtotal CSS; B) verbal IQ standard subscore; C) SWAN inattention index average score. Each dot on the scatter plot indicates a child's score. The violin plots model the distribution of the scores. The boxplots show the quartile ranges of the data. Data of children passing a specified scan are colored in blue; those failing or not attempting (NA) that scan a illustrated are colored orange.  Abbreviations: R-fMRI, first resting state fMRI; T-fMRI, task fMRI; R-fMRI 2, second resting state fMRI; DTI, diffusion tensor imaging; SWAN, Strengths and Weaknesses of Attention-Deficit/Hyperactivity Disorder Symptoms and Normal Behavior Scale; ADOS-2 CSS RRB, Autism Diagnosis Observation Schedule Calibrated Severity Score Restricted Repetitive Behaviors.

**Supplemental Figure 3**

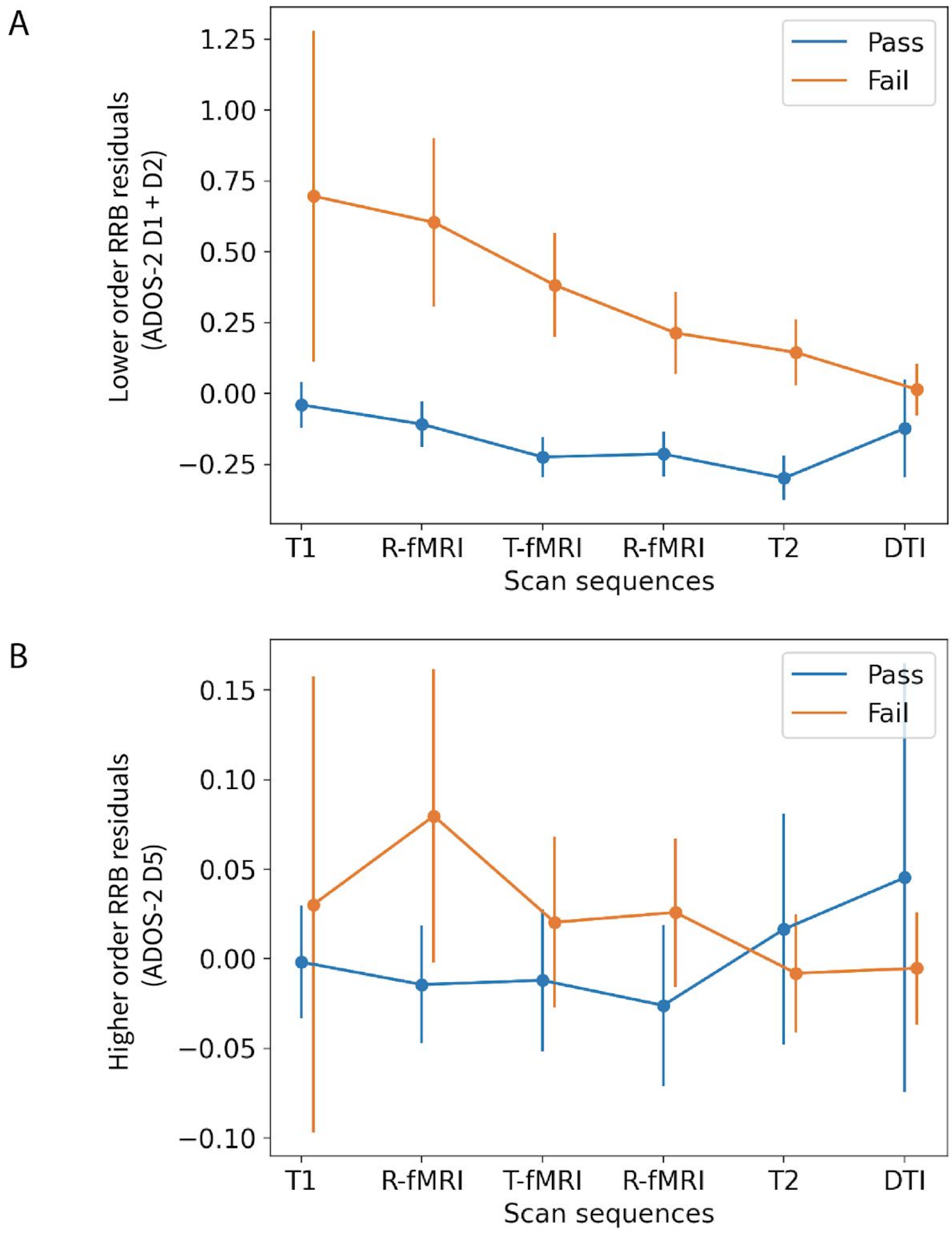


**Supplemental Figure 3 - Age-Residualized ADOS-2 lower- and higher-order RRB item scores.** Group mean and standard errors for A) lower (D1+D2 item) and B) higher order RRB item (D5) tafter regressing out for age (the strongest predictive feature) for the children failing (orange) and passing Q/A (blue) for each of the scans in the session. The regression models were as follows: Higher order ~ Age + e, and Lower order ~ Age + e, where e is the error. Consistent with (Bishop et al. 2013) lower and higher order item scores can be derived for ADOS-2 module 3, as such these scores are from children completing ADOS-2 module 3 exam (N = 184 out of 201; 92%).

**Supplemental Figure 4**

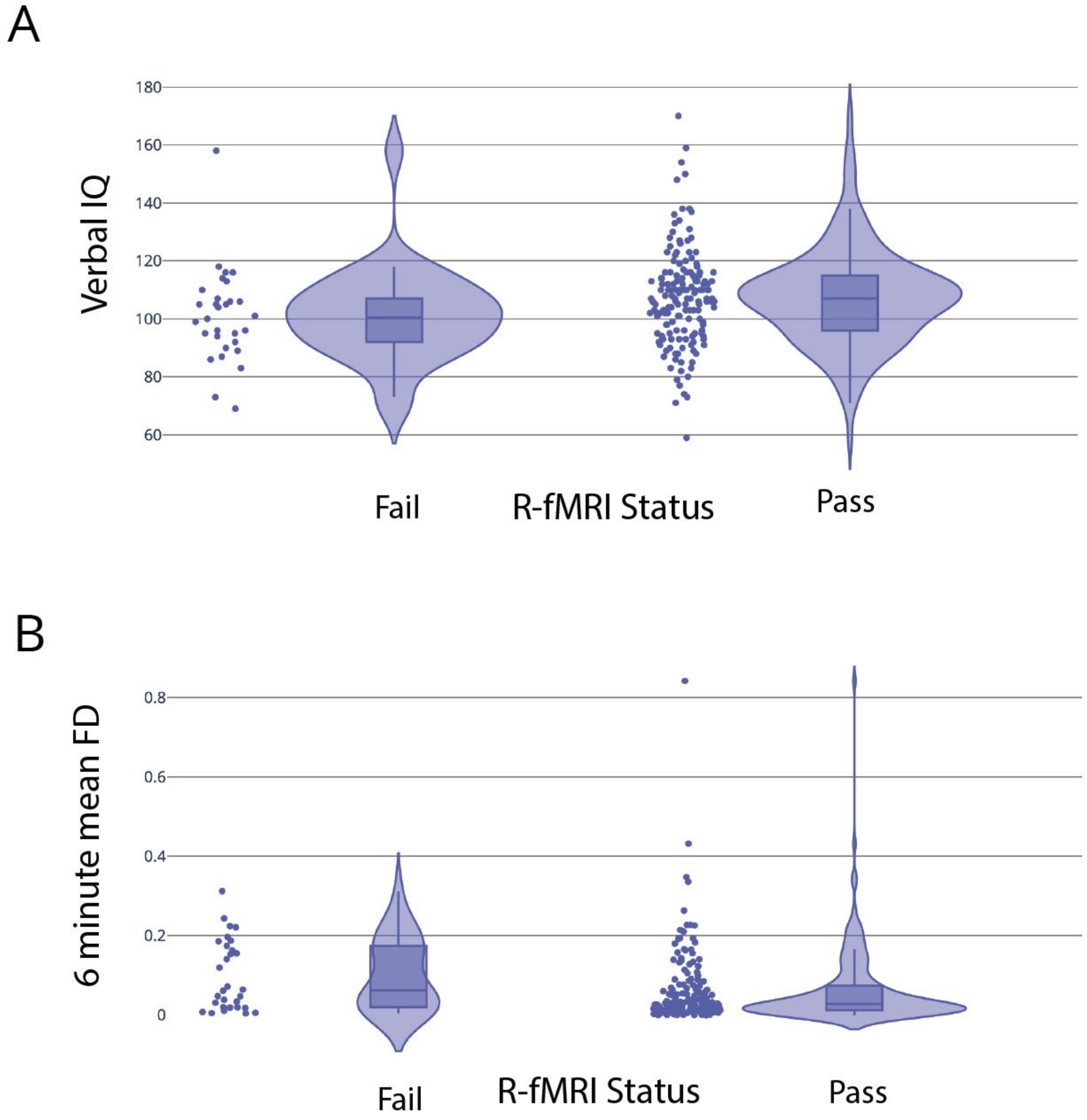


**Supplemental Figure 4 - Secondary predictive features associated with the minimal set of scan success.** Scatter, violin, and box plots showing the group distribution of A) verbal IQ standard scores and B) mean framewise displacement (FD) of the last 6 min step of the simulator training for the children pasing and those failing the minimal scan set (T1-weighted + R-fMRI1). Each dot on the scatter plot indicates a child's score. The violin plots model the distribution of the scores. The boxplots show the quartile ranges of the data.

**Supplemental figure 5**

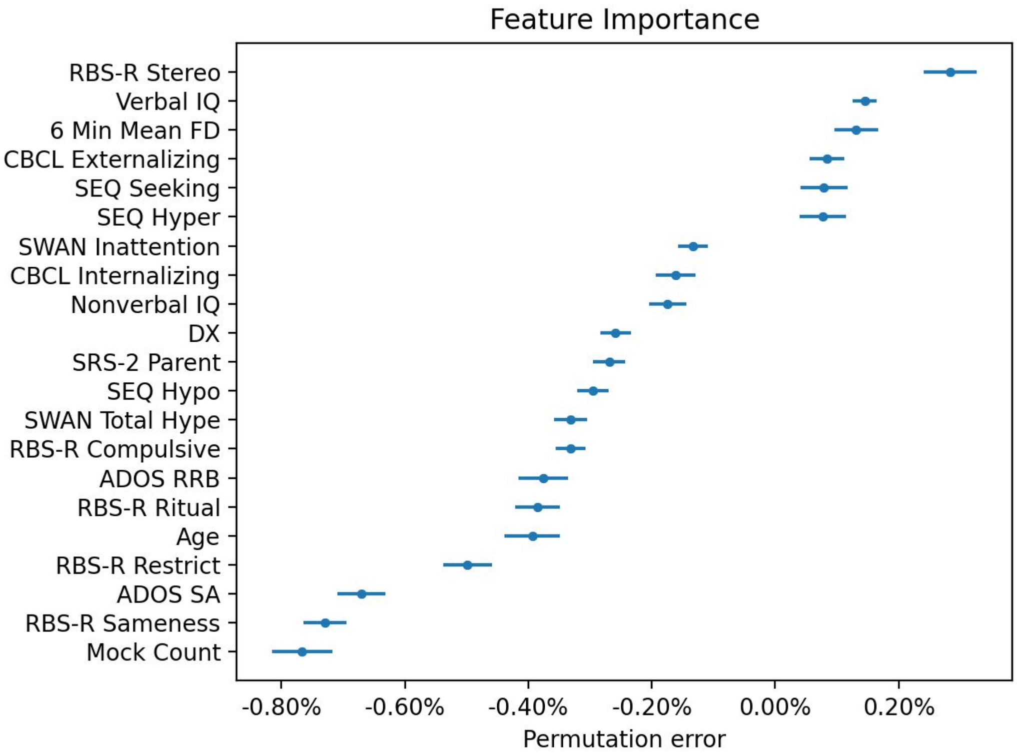


**Supplemental Figure 5. Feature importance after adding diagnosis (DX) to the naive bayes classification model.** Feature importance after adding diagnostic label ($ADHD_{w/oASD}$ vs ASD) as a feature to the naive bayes classification model. After adding diagnosis as a feature, the model’s virtually did not change (accuracy= 0.74, precision= 0.84 and recall=0.86).

**Supplementary Table 1**

| Variable | NYU (N=133) | | CMI (N=117) | | | In-between group differences | |
|---|---|---|---|---|---|---|---|
| | Mean | SD | Mean | SD | *df* | *W* | p-value (FDR corrected) |
| Age, (years) | 8.46 | 1.75 | 8.75 | 1.69 | 248 | 6919.5 | 0.31 |
| IQ (standard score) | | | | | | | |
| Full scale IQ | 100 | 15 | 104 | 18 | 248 | 6966.0 | 0.31 |
| Non verbal IQ | 102 | 16 | 106 | 18 | 248 | 6721.0 | 0.25 |
| Verbal IQ | 99 | 17 | 104 | 18 | 248 | 6379.0 | 0.20 |
| ADOS-2 CSS | | | | | | | |
| Total | 5.17 | 2.76 | 4.75 | 2.89 | 248 | 7032.0 | 0.33 |
| RRB | 4.67 | 3.00 | 5.02 | 3.30 | 248 | 7264.5 | 0.40 |
| Social affect | 5.68 | 2.54 | 5.09 | 2.67 | 248 | 6756.5 | 0.25 |
| Child behavior checklist T scores | | | | | | | |
| Total problems | 61.77 | 9.06 | 62.92 | 9.17 | 237 | 6649.0 | 0.40 |
| Externalizing problems | 58.36 | 10.08 | 59.19 | 10.79 | 238 | 6790.0 | 0.46 |
| Internalizing problems | 57.96 | 10.58 | 59.49 | 11.06 | 239 | 6654.0 | 0.40 |
| RBS-R Repetitive Behavior Scale raw scores | | | | | | | |
| Total | 15.16 | 13.27 | 18.39 | 17.92 | 240 | 6989.5 | 0.40 |
| Compulsive | 1.82 | 2.44 | 2.34 | 3.54 | 238 | 6922.5 | 0.40 |
| Restrictive | 1.76 | 2.06 | 2.04 | 2.53 | 238 | 7177.0 | 0.50 |
| Ritualistic | 3.04 | 3.18 | 3.44 | 3.69 | 238 | 7166.0 | 0.50 |
| Sameness | 4.83 | 4.63 | 5.49 | 5.58 | 238 | 7081.5 | 0.49 |
| Stereotypical | 2.67 | 2.72 | 2.99 | 3.62 | 238 | 7261.5 | 0.50 |
| SEQ | | | | | | | |
| Hyperresponsiveness | 28.45 | 7.68 | 28.45 | 7.53 | 226 | 6382.0 | 0.50 |
| Hyporesponsiveness | 10.72 | 2.76 | 11.59 | 3.57 | 226 | 5700.5 | 0.31 |
| Seeking | 25.57 | 8.80 | 26.39 | 8.01 | 226 | 6026.5 | 0.40 |
| SRS-2 parent T score | | | | | | | |
| Total | 62.33 | 11.05 | 62.41 | 12.21 | 239 | 7152.5 | 0.52 |
| SWAN average scores | | | | | | | |
| Total | 1.01 | 0.79 | 1.05 | 0.85 | 240 | 7120.5 | 0.50 |
| Hyperactivity/impulsivity | 0.93 | 0.91 | 0.97 | 0.98 | 240 | 7172.5 | 0.50 |
| Inattention | 1.10 | 0.92 | 1.12 | 0.94 | 240 | 7198.5 | 0.50 |
| | # | % | # | % | df | $\chi 2$ | p-value (FDR corrected) |
| Male | 106 | 73 | 93 | 86 | 1 | 5.08 | 0.91 |
| Primary diagnosis (#ASD) | 61 | 46 | 51 | 44 | 1 | 0.05 | 0.85 |
| Socioeconomic status class (Class 4, 5 versus 1, 2, 3) | 95 | 71 | 76 | 65 | 1 | 1.09 | 0.49 |
| Presence psychiatric comorbidities (yes) | 72 | 54 | 81 | 69 | 1 | 5.35 | 0.25 |
| Medication status (naïve) | 93 | 70 | 79 | 68 | 1 | 0.20 | 0.71 |

**Table 1. Characteristics of subjects by site.** ***Notes and Abbreviations***: ADHD, attention-deficit/hyperactivity disorder; ADOS-2, Autism Diagnostic Observation Schedule, second edition; ASD, autism spectrum disorder; CSS, calibrated severity scores; SA, social affect; RRB, restricted and repetitive behaviors; SCQ, Social Communication Questionnaire, SRS-P, Social Responsiveness Scale by Parents; SWAN, Strengths and Weaknesses of Attention-Deficit/Hyperactivity

symptoms and Normal behaviors; SEQ, Sensory Experience Questionnaire. For between-group comparisons, Mann-Whitney U tests were computed for continuous variables. Chi-squared tests were calculated for categorical variables. Children who were missing two or fewer instruments had imputed values, as described in the methods section below and in the main manuscript. Detailed information about the sample is presented in Table 2 of the main manuscript.

**Supplemental Table 2**

| | Matrix | Sl # | FOV | FOV Phase % | Res (mm) | TR ms | TE ms | TI ms | FA o | MBA | PPF | dirs | b | TA min:sec | Vol # |
|---|---|---|---|---|---|---|---|---|---|---|---|---|---|---|---|
| ***T1-w*** | 320x300 | 208 | 256x256 | 93.8 | 0.8x0.8 x0.8 | 240 0 | 2.2 4 | 10 60 | 8 | - | off | - | - | 6:36 | - |
| ***R1*** | 90x92 | 66 | 216x216 | 102.2 | 2.4x2.4 x2.4 | 800 | 30 | - | 55 | 6 | off | - | - | 6:20 | 465 |
| ***Task 1*** | 90x92 | 66 | 216x216 | 102.2 | 2.4x2.4 x2.4 | 800 | 30 | - | 55 | 6 | off | - | - | 4:39 | 339 |
| ***Task 2*** | 90x92 | 66 | 216x216 | 102.2 | 2.4x2.4 x2.4 | 800 | 30 | - | 55 | 6 | off | - | - | 4:39 | 339 |
| ***R2*** | 90x92 | 66 | 216x216 | 102.2 | 2.4x2.4 x2.4 | 800 | 30 | - | 55 | 6 | off | - | - | 4:39 | 339 |
| ***T2-w*** | 320x300 | 208 | 256x256 | 93.8 | 0.8x0.8 x0.8 | 320 0 | 564 | - | - | - | on | - | - | 5:57 | - |
| ***DTI*** | 140x140 | 81 | 240x240 | 100 | 1.7x1.7 x1.7 | 453 0 | 89. 6 | - | 90 | - | 6/8 | 137 | 3000 | 10:43 | - |

**Supplemental Table 2 — *MRI imaging parameters at each scan in the protocol.*** Acronyms used: b, b-values; dirs, number of diffusion directions; DTI, diffusion tensor imaging; FA, flip angle; FOV, field of view; MBA, multiband acceleration; TA, time of acquisition; TE, echo time; TI, inversion time; TR, repetition time; T1-w, T1-weighted (MPRAGE); T2-w, T2 weighted; PPF, phase partial Fourier; Res; spatial resolution; R1, R-fMRI 1st scan; R2, R-fMRI 2nd scan; task 1, task-fMRI 1st scan block; Sl; slice #; task 2, task-fMRI 2nd scan block; Vol, number of volumes,

**Supplemental Table 3 Characteristic of the enrolled sample by primary diagnosis**

| Variable | ADHD (N=138)[m] | | ASD (N=112) | | | In-between group differences | |
|---|---|---|---|---|---|---|---|
| | Mean | SD | Mean | SD | *df* | *W* | p-value |
| Age, (years) | 8.8 | 1.7 | 8.7 | 1.9 | 248 | 8094 | 0.520 |
| IQ[a] | | | | | | | |
| Full scale IQ | 104 | 15 | 99 | 18 | 248 | 9101 | 0.016 |
| Non verbal IQ | 103 | 18 | 100 | 18 | 247 | 8379 | 0.211 |
| Verbal IQ | 107 | 16 | 101 | 18 | 248 | 9285 | 0.006 |
| ADOS-2[b] CSS | | | | | | | |
| Total | 3.7 | 2.2 | 7.1 | 2.3 | 248 | 2359 | $p<0.0001$ |
| RRB | 3.5 | 2.7 | 6.9 | 2.9 | 248 | 3034 | $p<0.0001$ |
| Social affect | 4.4 | 2.3 | 7.1 | 2.1 | 248 | 3012 | $p<0.0001$ |
| Child behavior checklist T scores[c] | | | | | | | |
| Total problems | 61.3 | 9.1 | 63.7 | 9.1 | 238 | 5984 | 0.039 |
| Externalizing problems | 59.2 | 10.4 | 58.2 | 10.5 | 238 | 6097 | 0.049 |
| Internalizing problems | 57.3 | 11.6 | 60.0 | 10.0 | 239 | 7468 | 0.476 |
| RBS-R Repetitive Behavior Scale raw scores[d] | | | | | | | |
| Total | 11.4 | 13.3 | 23.0 | 16.3 | 240 | 3461 | $p<0.0001$ |
| Compulsive | 1.5 | 3.0 | 2.8 | 3.0 | 238 | 4526 | 0.001 |
| Restrictive | 1.1 | 1.8 | 2.9 | 2.5 | 238 | 3402 | $p<0.0001$ |
| Ritualistic | 2.0 | 2.8 | 4.6 | 3.6 | 238 | 3569 | $p<0.0001$ |
| Sameness | 3.8 | 4.6 | 6.7 | 5.3 | 238 | 4327 | $p<0.0001$ |
| Self-injurious | 1.1 | 1.8 | 1.9 | 2.3 | 238 | 5737 | 0.005 |
| Stereotypical | 1.9 | 2.3 | 4.0 | 3.4 | 238 | 4265 | $p<0.0001$ |
| SCQ lifetime raw score[e] | | | | | | | |
| Total | 5.9 | 5.3 | 14.6 | 6.4 | 218 | 1671 | $p<0.0001$ |
| SEQ[f] | | | | | | | |
| Hyperresponsiveness | 25.9 | 7.7 | 31.8 | 7.0 | 201 | 2584 | $p<0.0001$ |
| Hyporesponsiveness | 10.3 | 3.0 | 12.1 | 3.5 | 201 | 3398 | $p<0.0001$ |
| Seeking | 23.4 | 8.6 | 29.3 | 8.0 | 201 | 2949 | $p<0.0001$ |
| Non-social | 40.0 | 11.6 | 50.1 | 10.6 | 201 | 2512 | $p<0.0001$ |
| Social | 18.2 | 5.033 | 21.2 | 4.8 | 201 | 3102 | $p<0.0001$ |
| SRS-2 parent T score[g] | | | | | | | |
| Total | 57.29 | 10.37 | 69.0 | 9.7 | 240 | 2935 | $p<0.0001$ |
| SWAN average scores[h] | | | | | | | |
| Total | 0.97 | 0.80 | 1.1 | 0.8 | 236 | 6306 | 0 |
| Hyperactivity/impulsivity | 0.86 | 0.92 | 1.1 | 1.0 | 236 | 6101 | 0 |
| Inattention | 1.09 | 0.94 | 1.1 | 0.9 | 236 | 6683 | 1 |
| | # | % | # | % | df | $\chi 2$ | p-value |
| Male | 101 | 73 | 96 | 86 | 1 | 5.081 | 0.024 |
| Hispanic[i] | 29 | 21 | 36 | 32 | 1 | 3.377 | 0.066 |
| Race[i] | | | | | 3 | 2.720 | 0.437 |
| White | 84 | 61 | 58 | 52 | | | |
| Black | 16 | 12 | 18 | 16 | | | |
| Mixed | 21 | 15 | 18 | 16 | | | |
| Other | 14 | 25 | 16 | 14 | | | |
| Socioeconomic status class (SES)[j] | | | | | 1 | 0.904 | 0.342 |
| Class 4 and 5 | 99 | 72 | 72 | 64 | | | |
| # Psychiatric comorbidities[k] | | | | | 2 | 4.733 | 0.094 |
| One | 43 | 31 | 49 | 44 | | | |
| Two | 13 | 9 | 34 | 30 | | | |
| Three or more | 6 | 4 | 8 | 7 | | | |
| Medication status[l] | | | | | 1 | 2.8367 | 0.092 |
| Medication naïve | 103 | 75 | 69 | 62 | | | |

**Table 3.** ***Notes and Abbreviations***: ADHD, attention-deficit/hyperactivity disorder; ADOS-2, Autism Diagnostic Observation Schedule, second edition; ASD, autism spectrum disorder; CSS, calibrated severity scores; RRB, restricted and repetitive behaviors; SA, social affect; SCQ, Social Communication Questionnaire; SES, socioeconomic status; SEQ, Sensory Experience Questionnaire; SRS-P, Social Responsiveness Scale by Parents; SWAN, Strengths and Weaknesses of Attention-Deficit/Hyperactivity symptoms and Normal behaviors; [a] One child with ADHD was not administered one non-verbal IQ subscale. [b] Most children were administered module 3 of the ADOS-2, except for one child with ADHD and 11 with ASD who were administered ADOS-2 module 2. [c]Parent T scores on CBCL total, internalizing and externalizing subscales were not collected for 7 children with ASD and for 3 children with ADHD. [d] Parent RBS-R scores were not available for 6 children with ADHD and 2 children with ASD. [e] Parent SCQ Lifetime Total scores were not collected for 16 children with ADHD and 13 children with ASD.[f] Parent SEQ assessments were not collected for 26 children with ASD and 21 children with ADHD. [g] Parent SRS-2 T Total scores were not available for 6 children with ASD and 2 children with ADHD.[h] SWAN questionnaires were not collected for 4 children with ADHD and 8 children with ASD. [i]Information on ethnicity and race was not available for 3 children with ADHD and 2 with ASD. “Other” includes American Indian/Alaskan Native, Native Hawaiian/Pacific Islander, mixed or other not specified. Of the 35 children with ADHD, 8 identified as Asian, 6 as other. Of the 33 children with ASD, 7 identified as Asian, 1 as American Indian/Alaskan Native, 1 as Native Hawaiian/other Pacific islander, and 7 as ‘other.’ [j] Data on SES was missing for 7 children with ASD and 6 children with ADHD. The ADHD group included n=4 children in SES Class 1, n=12 children in SES Class 2, n=17 in SES Class 3, n=31 in SES Class 4 and n=68 SES Class 5. The ASD group included: n=6 children in SES Class 1, n=9 in SES Class 2, n=18 in SES Class 3, n=27 in SES Class 4, and n=44 in SES Class 5. [k] Information on DSM-5 categorical comorbidities was not available for 1 child with ASD. [l] Information on medication status was not available for 4 children with ASD. For the 35 children with ADHD who were not medication naïve, 24 were on current medication and 11 were off medication but not naïve. For the 39 children with ASD who were not medication naïve, 28 were on current medication and 11 were off medication but not naïve. [m] Of 138 children with ADHD, n=72 as combined, n=46 were classified as predominantly inattentive, n=6 as predominantly hyperactive/impulsive, and 14 as ADHD otherwise specified.
